# BAMM-SC: A Bayesian mixture model for clustering droplet-based single cell transcriptomic data from population studies

**DOI:** 10.1101/392662

**Authors:** Zhe Sun, Li Chen, Hongyi Xin, Qianhui Huang, Anthony R Cillo, Tracy Tabib, Ying Ding, Jay K Kolls, Tullia C Bruno, Robert Lafyatis, Dario AA Vignali, Kong Chen, Ming Hu, Wei Chen

## Abstract

The recently developed droplet-based single cell transcriptome sequencing (scRNA-seq) technology makes it feasible to perform a population-scale scRNA-seq study, in which the transcriptome is measured for tens of thousands of single cells from multiple individuals. Despite the advances of many clustering methods, there are few tailored methods for population-scale scRNA-seq studies. Here, we have developed a **B**Ayesiany **M**ixture **M**odel for **S**ingle **C**ell sequencing (BAMM-SC) method to cluster scRNA-seq data from multiple individuals simultaneously. Specifically, BAMM-SC takes raw data as input and can account for data heterogeneity and batch effect among multiple individuals in a unified Bayesian hierarchical model framework. Results from extensive simulations and application of BAMM-SC to in-house scRNA-seq datasets using blood, lung and skin cells from humans or mice demonstrated that BAMM-SC outperformed existing clustering methods with improved clustering accuracy and reduced impact from batch effects. BAMM-SC has been implemented in a user-friendly R package with a detailed tutorial available on www.pitt.edu/~Cwec47/singlecell.html.

## 1. Introduction

Single cell RNA sequencing (scRNA-seq) technologies have been widely used to measure gene expression for each individual cell, facilitating a deeper understanding of cell heterogeneity and better characterization of rare cell types (Gawad, et al., 2016; Tang, et al., 2009). Compared to early generation scRNA-seq technologies, the recently developed droplet-based technology, largely represented by the 10X Genomics Chromium system, has quickly gained popularity because of its high-throughput (tens of thousands of single cells per run), high efficiency (a couple of days), and relatively lower cost (< $2 per cell) (Macosko, et al., 2015; Zheng, et al., 2017; Jaitin, et al., 2014; Pollen, et al., 2014). It is now feasible to conduct population-scale single cell transcriptomic profiling studies, where several to tens or even hundreds of individuals are sequenced (van der Wijst, et al., 2018).

A major task of analyzing droplet-based scRNA-seq data is to identify clusters of single cells with a similar transcriptomic profile. To achieve this goal, unsupervised clustering methods such as K-means clustering, hierarchical clustering, and density-based clustering approaches (Rodriguez, et al., 2014) have been widely used. Recently, single cell interpretation via multikernel learning (SIMLR) (Wang, et al., 2017), CellTree (duVerle, et al., 2016), SC3 (Kiselev, et al., 2017) and DIMM-SC (Sun, et al., 2017) have been proposed for clustering droplet-based scRNA-seq data. However, it is still unclear how these methods can be scaled up to study population-scale droplet-based scRNA-seq data with tens of thousands of cells collected from many individuals and conditions. In addition, no existing clustering method is tailored for modeling multiple layers of heterogeneity imbedded in population studies. In the typical analysis of population-scale scRNA-seq data, reads from each individual are processed separately and then merged together for downstream analysis. For example, in 10X Genomics CellRanger pipeline, to aggregate multiple libraries, reads from different libraries are down-sampled such that all libraries have the same sequencing depth, leading to substantial information loss for individuals with higher sequencing depth. Alternatively, reads can be naively merged across all individuals without any library adjustment, leading to batch effect and unreliable clustering results. Although heterogeneity among multiple individuals can be reduced using the recently developed method implemented in Seurat (Satija, et al., 2015) and scran (Lun, et al., 2016), these methods require raw counts to be transformed to continuous numbers, leading to interpretation difficulty and potential over-correction.

We first conducted an exploratory data analysis to demonstrate the existence of batch effect in multiple individuals using both public and in-house droplet-based scRNA-seq datasets from human. We isolated peripheral blood mononuclear cells (PBMCs) from whole blood obtained from 4 healthy donors and used the 10X Chromium system to generate scRNA-seq data. We also included one additional healthy donor from a published PBMC scRNA-seq data (Zheng, et al., 2017) to increase the complexity of the testing dataset. Detailed sample information was summarized in Fig. 1a and Supplementary Table 1. In this cohort, sample 1 and sample 2 were sequenced in one batch; sample 3 and sample 4 were sequenced in another batch; sample 5 was downloaded from the original study conducted by 10X Genomics (Zheng, et al., 2017). As an exploratory data analysis, we made a t-SNE plot based on the first 50 PCs (Supplementary Fig. 1) of these 5 donors, and observed a clear batch effect: samples from the same batch tend to cluster together. This illustrative example demonstrates the importance and urgent needs of charactering different levels of variability and removing potential batch effect among droplet-based scRNA-seq datasets collected from multiple individuals. In the following sections, we will briefly describe our method, evaluate its performance in simulation studies, and apply it to three local scRNA-seq datasets including PMBC, skin, and lung tissues from human or mouse.

**Figure 1.**
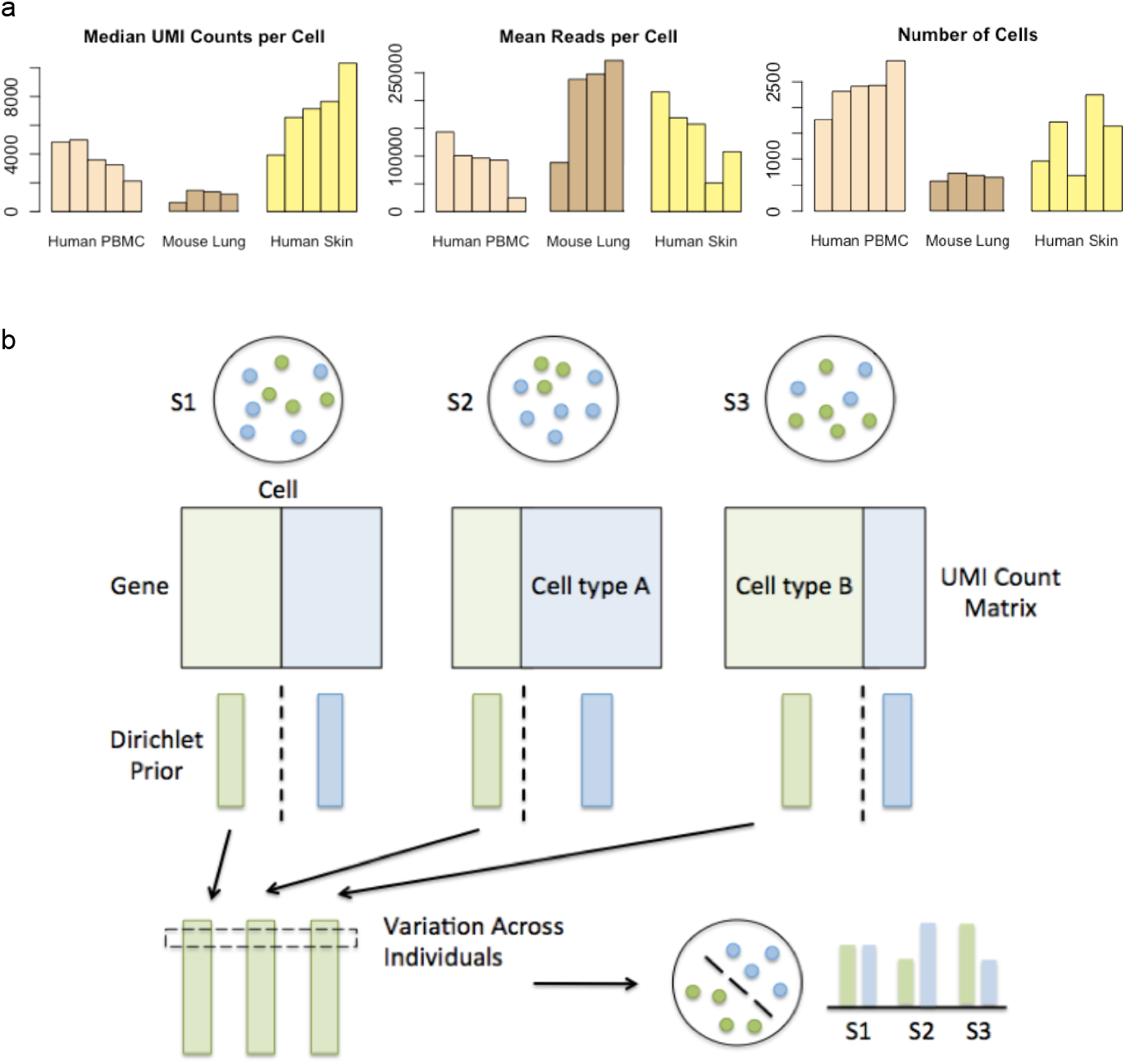
(a) Sample information of three droplet-based scRNA-seq datasets. (b) Overview of clustering with BAMM-SC.

## 2. Methods

### 2.1 Statistical model

We propose a Bayesian hierarchical Dirichlet multinomial mixture model to explicitly characterize different levels of variability in population scale scRNA-seq data. Specifically, let *x_iji_* represent the number of unique UMIs for gene *i* in cell *j* from individual *l* (1 ≤ *i* ≤ *G*, 1 ≤ *j* ≤ *C_l_*, 1 ≤ *l* ≤ *L*). Here *G, C_l_* and *L* denote the total number of genes, cells in individual *l* and individuals, respectively. Our goal is to perform simultaneously clustering for all *L* individuals. We assume that among each individual, all single cells consist of *K* distinct cell types. Here *K* is pre-defined according to prior biological knowledge, or will be estimated from the model, and *K* is the same among all *L* individuals.

Assume that ***x*_1*jl*_**, *x*_2*jl*_, … *x_Gjl_*, the gene expression for cell *j* in individual *l*, follows a multinomial distribution *Multi*(*T_jl_*, ***p_.jl_***). Here 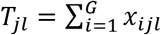 is the total number of UMIs, (***p_.jl_*** = (*p*_1*jl*_, *p*_2*jl*_, …, *p_Gjl_* is the gene expression vector. In addition, let *z_jl_* ∊ {1,2,…, *K*} represent the cell type label for cell *j* in individual *l*, where *z_jl_* = *k* indicates that cell *j* in individual *l* belongs to cell type *k*. Given *z_jl_* = *k*, we assume that ***p*_·jl_**; follows a Dirichlet prior *DIR*(**α**_**·l**(***k***)_), where ***α***_***·l***(***k***)_) = (*α*_1*l*(*k*)_, *α*_2*l*(*k*)_,…, *α*_*Gl*(*k*)_ (is the Dirichlet prior parameter for cell type *k* in individual *l*. Taken together, after integrating ***p_·jl_*** out, we have:

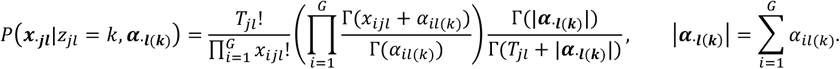

The joint distribution of ***x_·jl_*** and *z_jl_* is:

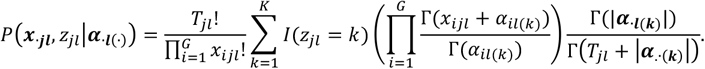

We then assume that all *C_l_* cells in individual *l* are all independent, then the joint distribution for individual *l* is as follows:

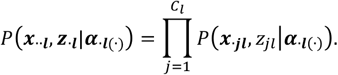

We further assume that all *L* individuals are independent, then the overall joint distribution is as follow

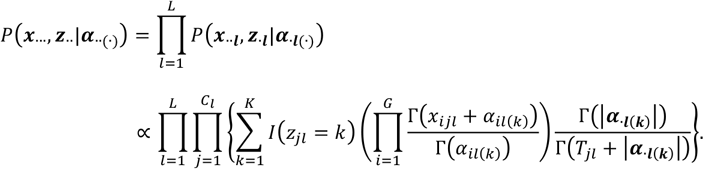

In this model, the two sets of parameters of interest are ***z..*** = {*Z_jl_* }_1*≤j≤C_*l*_,≤L*_’ the cell type label for cell *j* in individual *l*, and ***α..***_(∙)_ = {*α*_*il*(*k*)_}_1≤*i*≤*G*,≤*l*≤*L*,1≤*k*≤*K*_’ the Dirichlet parameters for gene *i* in cell type *k* in individual *l*. We adopt a full Bayesian approach and use Gibbs sample for the statistical inference. Specifically, the joint posterior distribution for ***z..*** and ***α..***_(∙)_ are:

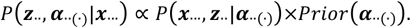

Since all *α*’s are strictly positive, we adopt a log-normal distribution as the prior distribution for *α*_il(*k*)_. We assume that for gene *i* in cell type *k*, *α*_*il*(*k*)_ from all *L* individuals share the same prior distribution 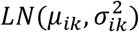).

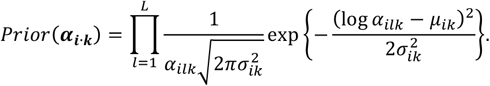

Here *μ_ik_* can be estimated by the mean of 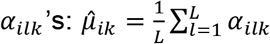. Estimation of 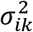 can be challenging due to limited number of individuals. We can assume all 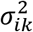’s follow a hyper-prior: Gamma distribution *Gamma*(*a_k_, b_k_*), to use information across all genes to estimate variance. In addition, we assume a non-informative prior for *μ_ik_*’s. Taken together, we have the full posterior distribution as follows:

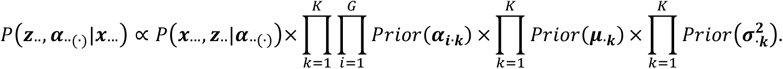

We use Gibbs sample to iteratively update *α*_*il*(*k*)_ and *z_ij_*. Details can be found in **Supplementary Material Section 1**.

### 2.2 Sequencing library construction

10X Genomics has recently released a commercialized droplet-based Chromium system, which is a microfluidics platform based on Gel bead in EMulsion (GEM) technology (Zheng et al., 2017). Cells mixed with reverse transcription reagents were loaded into the Chromium instrument. This instrument separated cells into mini-reaction “partitions” formed by oil microdroplets, each containing a gel bead and a cell, known as GEMs. GEMs contain a gel bead, scaffold for an oligonucleotide that is composed of an oligo-dT section for priming reverse transcription, and barcodes for each cell and each transcript as described (Zheng et al., 2017). GEM generation takes place in a multiple-channel microfluidic chip that encapsulates single gel beads. Reverse transcription takes place inside each droplet. Approximately 1,000-fold excess of partitions compared to cells ensured low capture of duplicate cells. The reaction mixture/emulsion was removed from the Chromium instrument, and reverse transcription was performed. The emulsion was then broken using a recovery agent, and following Dynabead and SPRI clean up cDNAs were amplified by PCR (C1000, Bio-Rad). cDNAs were sheared (Covaris) into ~200 bp length. DNA fragment ends were repaired, A-tailed and adaptors ligated. The library was quantified using KAPA Universal Library Quantification Kit KK4824 and further characterized for cDNA length on a Bioanalyzer using a High Sensitivity DNA kit. All sequencing experiments were conducted using Illumina NextSeq 500 in the Genomics Sequencing Core at the University of Pittsburgh.

### 2.3 Data description

#### 2.3.1 Human PBMC dataset

Peripheral blood was obtained from healthy donors by venipuncture. Peripheral blood mononuclear cells (PBMC) were isolated from whole blood by density gradient centrifugation using Ficoll-Hypaque. PBMC were then counted and resuspended in phosphate buffered saline with 0.04% bovinue serum albumin, and were processed through the Chromium 10X Controller according to the manufacturers’ instructions, targeting a recovery of ~2,000 cells. The following steps were all performed under the aforementioned protocol developed by 10X Genomics.

#### 2.3.2 Human Human Skin dataset

Skin samples were obtained by performing 3 mm punch biopsies from the dorsal mid-forearm of healthy control subjects after informed consent under a protocol approved by the University of Pittsburgh Institutional Review Board. Skin for scRNA-seq was digested enzymatically (Miltenyi Biotec Whole Skin Dissociation Kit, human) for 2 hours and further dispersed using the Miltenyi gentleMACS Octo Dissociator. The resulting cell suspension was filtered through 70 micron cell strainers twice and re-suspended in PBS containing 0.04% BSA. Cells from biopsies were mixed with reverse transcription reagents then loaded into the Chromium instrument (10X Genomics). ~2,600-4,300 cells were loaded into the instrument to obtain data on ~1,100-1,800 cells, anticipating a multiplet rate of ~1.2% of partitions. The following steps were all performed under the aforementioned protocol developed by 10X Genomics.

#### 2.3.3 Human Mouse lung dataset

Lung single cell suspension from naïve and infected C57BL/6 mice were subject to scRNA-seq library preparation protocol. Briefly, left lobs of both naïve and infected mice were removed and digested by Collagenase/DNase to obtain single cell suspension. Mononuclear cells after filtration with a 40μM cell strainer were separated into mini-reaction “partitions” or GEMs formed by oil micro-droplets, each containing a gel bead and a cell, by the Chromium instrument (10X Genomics). The reaction mixture/emulsion with captured and barcoded mRNAs were removed from the Chromium instrument followed by reverse transcription. The cDNA samples were fragmented and amplified using the Nextera XT kit (Illumina). The following steps were all performed under aforementioned the protocol developed by 10X Genomics.

## 3. Results

In this study, we propose a **BA**yesian **M**ixture **M**odel for **S**ingle **C**ell sequencing (BAMM-SC) to simultaneously cluster droplet-based scRNA-seq data from multiple individuals. BAMM-SC directly works on the raw counts without any data transformation and models the heterogeneity internally by learning variation of signature genes in a Bayesian hierarchical model framework.

Specifically, BAMM-SC adopts a Bayesian hierarchical Dirichlet multinomial mixture model, which explicitly characterizes three levels of variabilities (i.e., genes, cell types and individuals) (Fig. 1b and Methods). Our method has the following three key assumptions. First, cell type clusters are discrete, and each cell belongs to one cell type exclusively. Second, heterogeneity exists among different individuals, but the variability of same cell type among different individuals is much lower than the variability of different cell types among the same individual. Third, cells of the same cell type share a similar gene expression pattern. Compared to other clustering methods which ignore individual level variability, BAMM-SC has the following three key advantages: (1) BAMM-SC accounts for data heterogeneity among multiple individuals, such as unbalanced sequencing depths and technical biases in library preparation, and reduces the false positives of detecting individual-specific cell types. (2) BAMM-SC borrows information across different individuals, leading to improved power for detecting individual-shared cell types and higher reproducibility as well as stability of clustering results. (3) BAMM-SC provides a statistical framework to quantify the clustering uncertainty for each cell in the form of posterior probability for each cell type.

We have conducted comprehensive simulation studies to benchmark the performance of BAMM-SC. Specifically, we simulated droplet-based scRNA-seq data collected from multiple individuals from the posited Bayesian hierarchal Dirichlet multinomial mixture model (Methods and **Supplementary Material Section 2**). We considered two experimental designs, including (1) different heterogeneities among multiple individuals; and (2) different sequencing depths among multiple individuals. We applied BAMM-SC to each synthetic dataset, and compared the inferred cell type label of each single cell with the ground truth, measured by adjusted Rand index (ARI) (Rand, 1971) and normalized mutual information (NMI) (**Supplementary Material Section 3**). As a comparison, we also applied K-means clustering and DIMM-SC to synthetic dataset in two modes: (1) in mode 1, we pooled single cells from different individuals together while ignoring each individual label, and then applied K-means clustering / DIMM-SC to the pooled data; (2) in mode 2, we first applied K-means clustering / DIMM-SC to each individual separately, and then matched the inferred cell type label empirically. Specifically, we first obtained the cell clusters of the first individual and used it as a reference. Then, we matched each cell cluster of the rest individuals with the corresponding reference cell cluster, which has the highest Pearson correlation with its average gene expression profile. We denoted the two methods in mode 1 as K-means (Pooled) and DIMM-SC (Pooled), respectively, and the two methods in mode 2 as K-means (Sep) and DIMM-SC (Sep), respectively. Noticeably, our previous work has shown that DIMM-SC outperforms Seruat and CellTree when clustering scRNA-seq data from a single individual; therefore, we did not include Seruat and CellTree in our simulation studies. We simulated 100 datasets and applied clustering methods on each of them to evaluate the variability of the clustering results.

In the first experimental design, we simulated scRNA-seq data with different heterogeneities among multiple individuals. In our posited hierarchical model, the log normal prior distribution 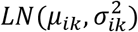 measures the heterogeneity of gene *i* in cell type *k* among multiple individuals. Without loss of generality, we used the mean of 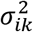 across all genes and all cell types to quantify the individual level heterogeneity. In the second experimental design, we simulated scRNA-seq data from multiple individuals with different sequencing depths. Specifically, we simulated data from 10 individuals and equally divided them into two groups, in which each group consists of 5 individuals. We set the sequencing depth of group 1 as half of the sequencing depth of group 2, and denoted these 10 individuals as the “Unequal” dataset. To avoid batch effect due to imbalanced sequencing depth, we down-sampled the sequencing depth of group 2 by half to match the sequencing depth of group 1, and denoted them as the “Equal” dataset. We applied BAMM-SC and the other four competing clustering methods, including K-means clustering (Pooled), DIMM-SC (Pooled), K-means clustering (Sep) and DIMM-SC (Sep), to both the “Unequal” and “Equal” datasets, and reported the corresponding ARIs and NMIs.

BAMM-SC consistently outperformed the other four competing methods across a variety of individual level heterogeneities by achieving higher average ARI and lower variation of ARI among 100 simulations as shown in Fig. 2a. As a comparison, K-means clustering and DIMM-SC are robust to different levels of individual heterogeneity. Noticeably, K-means-Sep and DIMM-SC-Sep provided xmore variable clustering results when *σ*^2^ is small, since it is sub-optimal to cluster each individual separately when individuals are relatively homogeneous. Fig. 2b shows that among all five clustering methods, BAMM-SC performed best by obtaining the highest average ARI and the lowest variation of ARI, for both “Unequal” and “Equal” datasets. Noticeably, K-means clustering performed worst for the “Unequal” dataset, as the unbalanced sequencing depths lead to systematic differences between individuals in group 1 and group 2. As a result, K-means clustering wrongly separated group 1 and group 2 as two distinct cell types, and failed to identify true cell types.

**Figure 2.**
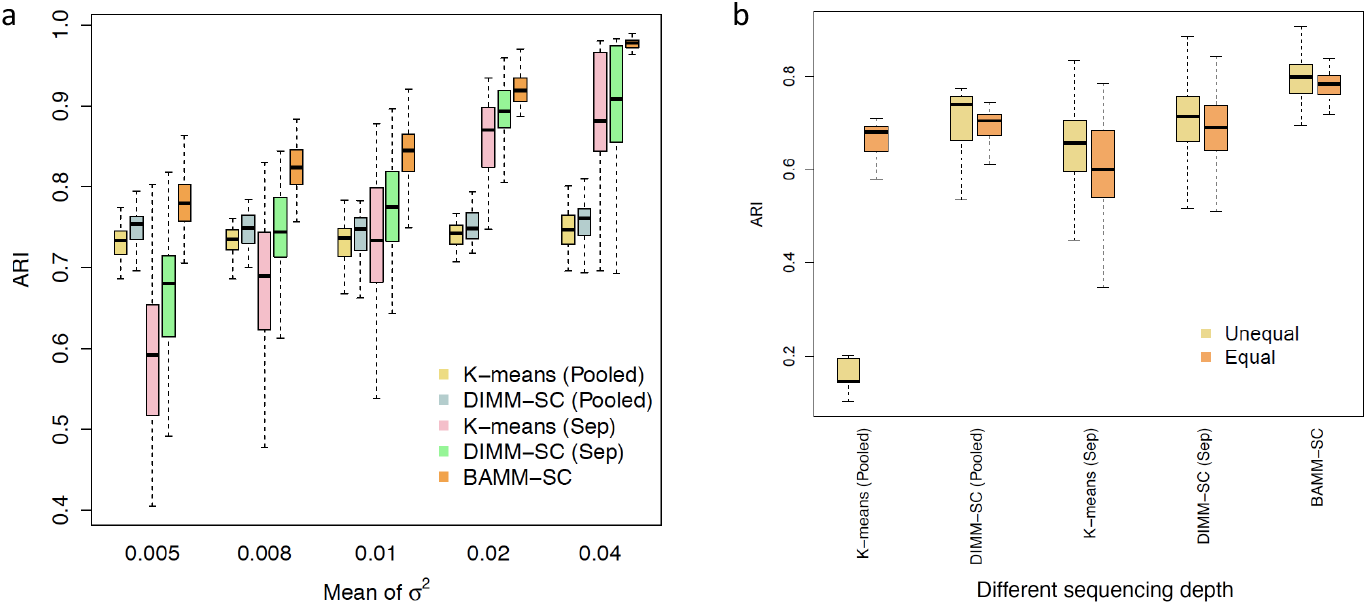
Boxplots of ARI for five clustering methods across 100 simulations, investigating how different heterogeneity among multiple individuals (a) and individuals with different sequencing depths (b) affect clustering results.

Furthermore, we performed comprehensive simulation analysis by generating simulated scRNA-seq datasets from different number of individuals (Supplementary Fig. 2a), different number of cell types (i.e., cluster) (Supplementary Fig. 2b), different overall sequencing depths (Supplementary Fig. 2c), different total number of cells (Supplementary Fig. 2d) and different cell-type-specific heterogeneities (i.e., the difference of gene expression profile between two distinct cell types) (Supplementary Fig. 2e), and obtained consistent results. BAMM-SC consistently outperformed other methods in terms of accuracy and robustness. We reached the same conclusion based on NMI (**Supplementary Fig. 3a – 3g**). Taken together, our comprehensive simulation studies have demonstrated that BAMM-SC is able to adaptively borrow information across multiple individuals, account for unbalanced sequencing depths, and provide more accurate and robust clustering results than other competing methods.

**Figure 3.**
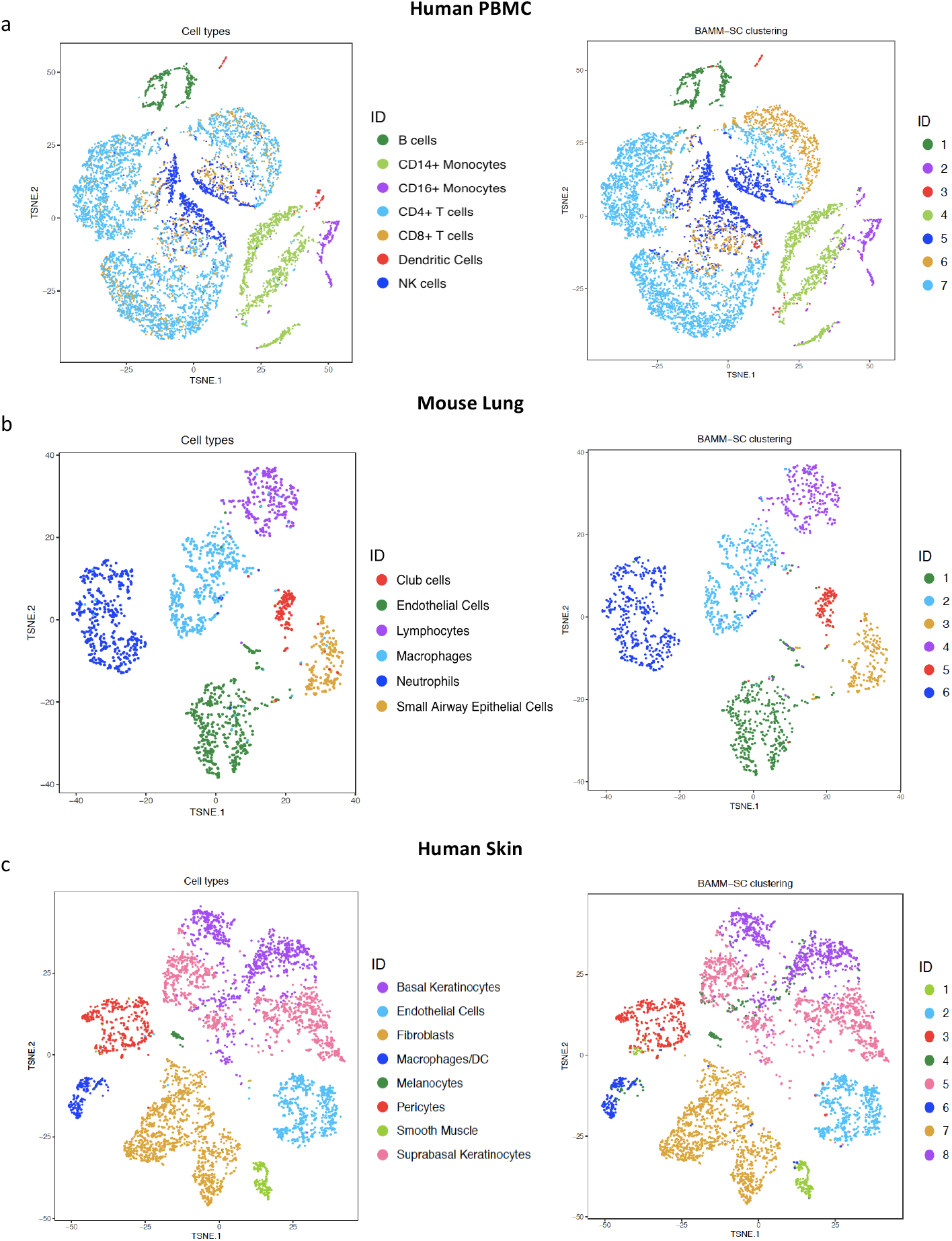
The t-SNE projection of cells from human PBMC (a), mouse lung (b) and human skin (c) tissues, colored by the approximated truth and BAMM-SC clustering results, separately.

To evaluate the performance of BAMM-SC in real applications, we further generated three scRNA-seq datasets including mouse lung, human PBMC and human skin tissues using 10X Chromium system at the University of Pittsburgh (see detailed data description in methods). The data are available for download (see data availability).

For aforementioned human PBMC samples, we first pooled cells from 5 donors together, filtered lowly expressed genes that were expressed in less than 1% cells. We then extracted the top 1, 000 highly variable genes based on their standard deviations. As shown in Supplementary Fig. 4, we identified 7 types of PBMCs based on the biological knowledge of cell-type-specific gene markers (Supplementary Table 2). Using these gene markers, >70% single cells can be assigned to a specific cell type. Since there is no gold standard for clustering analysis in this real dataset, we used the labels of these cells as the approximated truth to benchmark the clustering performance for different clustering methods. Cells with uncertain cell types are removed when calculating ARIs and NMIs.

We applied five clustering methods on these samples and repeated each method 10 times to evaluate the stability of its performance. In addition to K-means clustering, SC3, DIMM-SC and BAMM-SC, we also used scran (Lun, et al., 2016) to remove the batch effect among 5 samples and then applied K-means clustering on the normalized data (scran+K-means). The total number of clusters was set as 7, matching the biological knowledge from cell-type-specific gene markers. Fig. 3a (left figure) and Fig. 3a (right figure) show the t-SNE plots with each cell colored by their classification based on specific gene markers and cluster labels inferred by BAMM-SC, respectively. Despite some dendritic cells were wrongly identified as CD16+ Monocytes, these two plots are similar to each other (ARI=0.535, NMI=0.582) suggesting that BAMM-SC performed very well in human PBMC samples.

Based on the clustering results from BAMM-SC, we calculated the averaged cell proportions of each cell type among 10 runs for each PBMC sample. Fig. 4a shows that the proportions inferred from BAMM-SC is close to the proportions from the approximated truth based on gene markers, suggesting that BAMM-SC can account for batch effect. As shown in Fig. 5a, BAMM-SC achieved the highest ARI and NMI among all clustering methods. scran + K-means clustering slightly outperformed K-means clustering in terms of accuracy. We also generated t-SNE projection plots colored by cluster labels inferred by K-means clustering (Supplementary Fig. 5a), scran + K-means clustering (Supplementary Fig. 5b), SC3 (Supplementary Fig. 5c), DIMM-SC (Supplementary Fig. 5d) and BAMM-SC (Supplementary Fig. 5e). Noticeably, K-means clustering performed the worst for the combined PBMC datasets.

**Figure 4.**
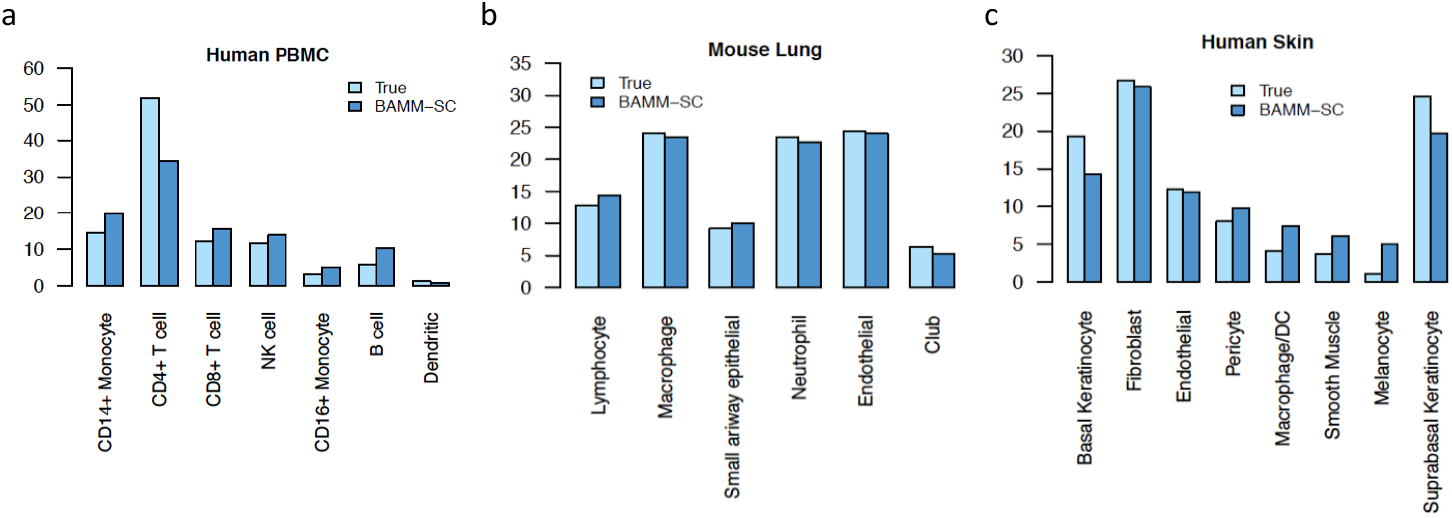
Bar plots of proportions of cell types among all individuals for human PBMC (a), mouse lung (b) and human skin (c) tissues, separately.

In addition to human PBMC samples, we also collected lung mononuclear cells from 4 mouse samples under 2 conditions: naive and streptococcus pneumonia (SP) infected. Among them, Sample 1 and Sample 2 are SP infected samples. Sample 3 and Sample 4 are naïve samples. Supplementary Fig. 6 shows the t-SNE plot of lung mononuclear cells from 4 mouse samples. Similar to the analysis of PBMC samples, after filtered lowly expressed genes, we pooled cells from 4 mice together and extracted the top 1,000 highly variable genes. As shown in Supplementary Fig. 7, we identified 6 types of cells based on the biological knowledge of cell-type specific gene markers (Supplementary Table 3). Taken together, >66% of single cells can be assigned to a specific cell type. Therefore, we used the labels of these cells as the approximated truth and removed cells with uncertain cell types from downstream analysis.

Fig. 3b **(left figure)** and Fig. 3b **(right figure)** shows the t-SNE plot with each cell colored by their cluster label based on cell-type-specific gene markers and cluster labels inferred by BAMM-SC, respectively. These two are highly similar (ARI=0.861, NMI=0.840), indicating the good performance of BAMM-SC. Fig. 5b **(left figure)** and Fig. 5b **(right figure)** shows that BAMM-SC outperformed other four clustering methods in terms of ARI and NMI. We also generated the t-SNE plots colored by cluster labels inferred by K-means clustering (Supplementary Fig. 8a), scran + K-means clustering (Supplementary Fig. 8b), SC3 (Supplementary Fig. 8c), DIMM-SC (Supplementary Fig. 8d) and BAMM-SC (Supplementary Fig. 8e). As shown in Supplementary Fig. 13, the proportions of neutrophils in SP infected samples (sample 1 and sample 2) are much higher than the proportions in naïve samples (sample 3 and sample 4). This is consistent with the fact that infections by bacteria and viruses may increase the number of neutrophils, which is a necessary reaction by the body (Chen, et al., 2013; Weiser, 2010). Interestingly, the proportion of cell types in naïve sample 3 is different from others, which may due to unsatisfied sample quality or unexpected bacterial infections.

**Figure 5.**
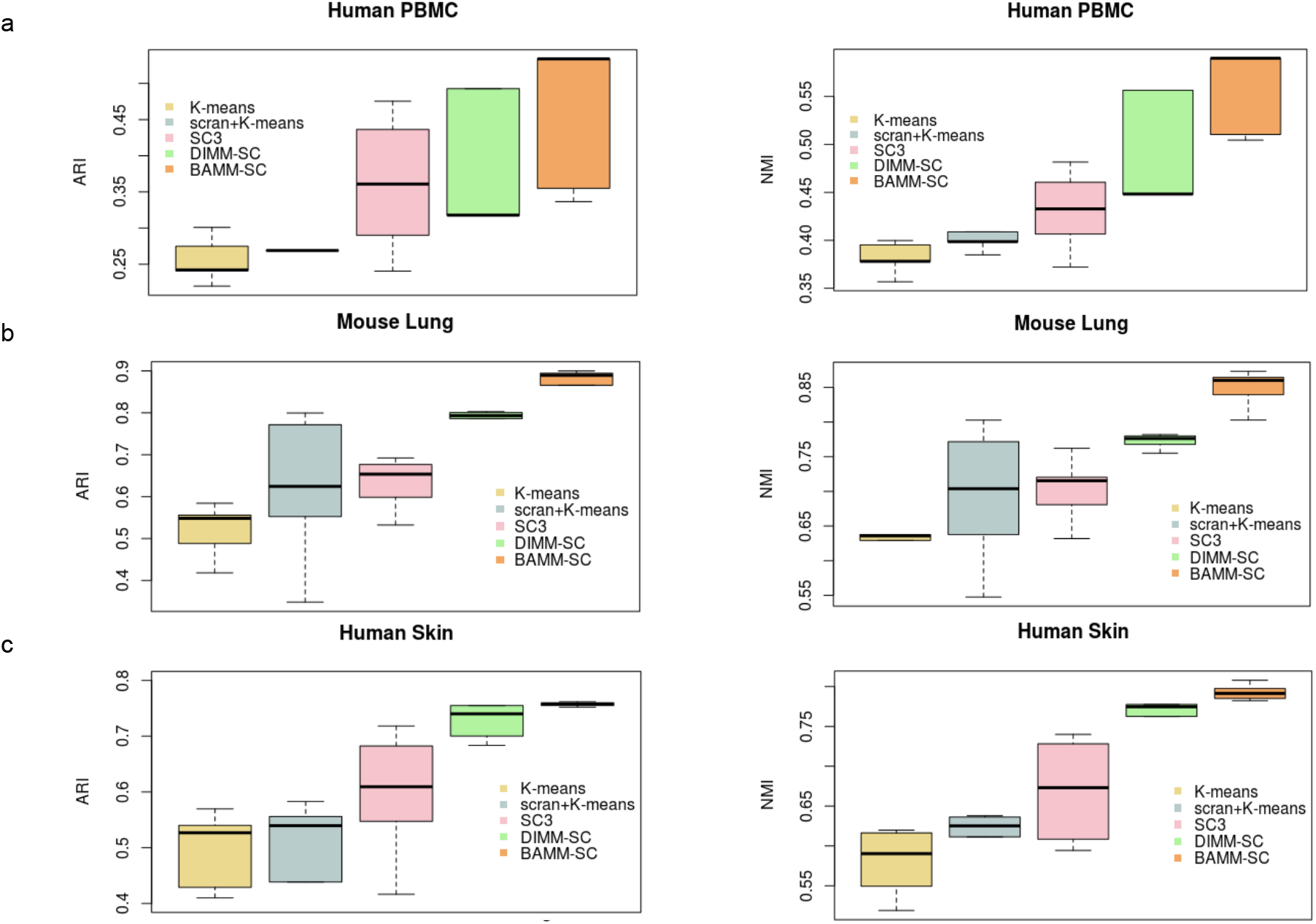
Boxplots of ARI and NMI from five clustering methods across 10 simulations for dataset human PBMC (a), mouse lung (b) and human skin (c), respectively.

To evaluate the clustering performance of BAMM-SC in solid tissue, we collected skin samples from 5 healthy donors that are part of a systemic sclerosis study (Tabib, et al., 2018). Fig. 1a lists the detailed sample information and Supplementary Fig. 9 shows the t-SNE plot of cells from 5 human skin samples. Similar to previous analyses, we first pooled cells from 5 samples together, removed lowly expressed genes, and then extracted the top 1,000 highly variable genes. As shown in Supplementary Fig. 10, we identified 8 major types of cells based on the biological knowledge of cell-type-specific gene markers (Supplementary Table 4). Taken together, >67% of single cells can be assigned to a specific cell type. We used the labels of these cells as the approximated truth and removed cells with uncertain cell types from downstream analysis.

As shown in Fig. 3c, BAMM-SC performed well in human skin samples, since the t-SNE plot with each cell colored by their cell type label based on gene markers is highly similar to the plot generated from the clustering result of BAMM-SC (ARI=0.786, NMI=0.810). Also, BAMM-SC achieved higher ARI and NMI compared with other clustering methods (t-test P-value < 0.01) (Fig. 5c). As comparisons, we generated the t-SNE plots colored by cluster labels inferred by K-means clustering (Supplementary Fig. 11a), scran + K-means clustering (Supplementary Fig. 11b), SC3 (Supplementary Fig. 11c), DIMM-SC (Supplementary Fig. 11d) and BAMM-SC (Supplementary Fig. 11e).

## 4. Software and data availability

BAMM-SC is available as an R package at http://www.pitt.edu/~wec47/BAMMSC.html. All the real data (human PBMC, mouse lung and human skin tissues) used in this study have been deposited at http://www.pitt.edu/~wec47/BAMMSC.html.

## 5. Discussion

In summary, we developed a novel Bayesian framework for clustering population-scale scRNA-seq data. BAMM-SC retains the raw data information by directly modeling UMI count without data transformation or normalization, facilitating straightforward biological interpretation. The Bayesian hierarchical model enables the joint characterization of multiple levels of uncertainty, including sampling variability, single cell level heterogeneity and individual level heterogeneity. Furthermore, BAMM-SC can borrow information across different individuals, leading to improved clustering accuracy. BAMM-SC is based on probabilistic models, thus providing the quantification of clustering uncertainty for each single cell.

Our model coupled with a computationally efficient MCMC algorithm is able to cluster large-scale droplet-based scRNA-seq data with feasible time cost. For example, it takes a couple hours to cluster over 10,000 cells from 5 individuals using the top 1,000 highly variable genes. Additionally, we can pre-define the number of clusters based on prior knowledge on the tissue or determine it using some model checking criteria such as AIC or BIC. As shown in Supplementary Fig. 12, AIC and BIC work as expected in the analysis of simulated datasets and provide a reliable range of cluster numbers for real data analysis based on prior knowledge. BAMM-SC is currently implemented in R/Rcpp with satisfactory computing efficiency to accommodate population scale scRNA-seq data. Further speed-up can be made through parallel computing.

There are several limitations of BAMM-SC. First, we filter out genes with excessive zeros from the analysis with the assumption that lowly-expressed genes do not contribute much to clustering. This may be a problematic for rare cell type identification. Second, we do not explicitly model a zero-inflation pattern, which may or may not affect clustering accuracy. A refined model needs to be further developed with a consideration of computational complexity and model flexibility. Third, in our model, we assume that each cell belongs to one distinct cluster. The posterior probability measures the clustering uncertainty, which cannot be interpreted as quantification of cell cycle or developmental stage. Our method has a potential to be extended to perform trajectory analysis (Trapnell, et al., 2014; Trapnell, 2015), which is beyond the scope of this paper.

We have applied BAMM-SC to three in-house datasets to showcase its feasibility of use with different tissue types and species. Consistent patterns of improvement were observed across different applications. We will test our method in many other datasets that are being rapidly generated at the University of Pittsburgh. With the increased popularity of population scale scRNA-seq studies, we expect that BAMM-SC will become a useful tool for elucidating single cell level transcriptomic heterogeneity.

## 6. Acknowledgements

This work is supported by National Institute of Health grants U54 DK107977 (M.H.), R01HG007358 (W.C.), R56HL137709 (K.C.), P50 CA097190 & P30 CA047904 (D.A.A.V.), R35HL139930 (J.K.) and Children’s Hospital of Pittsburgh (W.C. and Z.S.).

## Supplementary Material

### Section 1 Details of Gibbs sample

Based on Bayes formula, we have the full posterior distribution as follows:

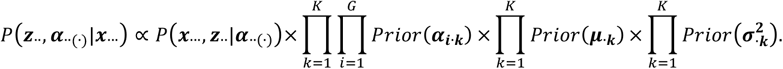

The complete log likelihood is:

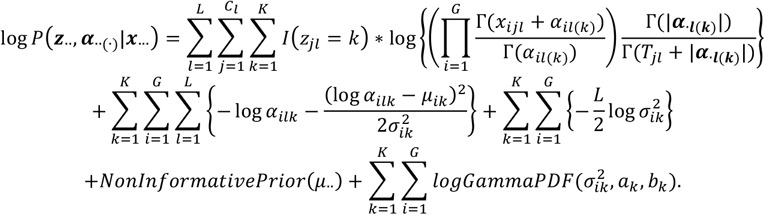

Here the hyper-prior parameters *a_k_* and *b_k_* can be pre-specified, or estimated from data via an empirical Bayes approach.

We will use Gibbs sample to iteratively update {*Z_jl_*}_1≤*j*≤*C_l_*,1≤,*l≤L*’_ {*α*_*il*(*k*)_}_1≤*i*≤*G*,≤*l*≤*L*,1≤*k*≤*K*_. For a given pair of *l* and *j*, the conditional distribution for *z_jl_* is a multinomial distribution, where

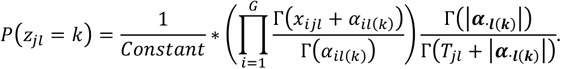

Where the normalization constant is:

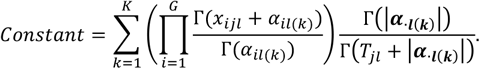

We will use random walk Metropolis within Gibbs to iteratively update *α*_*il*(*k*)_. For a given triple of *i*, *l* and *k*, the conditional log likelihood for *α*_*il*(*k*)_ is:

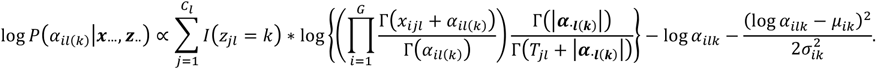

Similarly, we will use random walk Metropolis within Gibbs to iteratively update 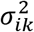. For a given pair of *i* and *k,* the conditional log likelihood for 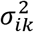 is:

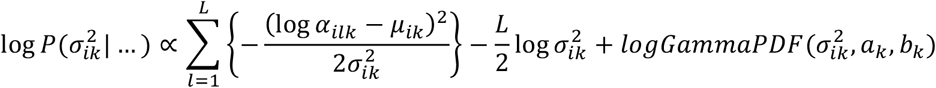

In random walk Metropolis algorithm, we adaptively select the step size of proposal distribution, to make sure that the acceptance rate is 20% ~ 30%.

### Label switching issue

We can first run DIMM-SC on all single cells pooled from all individuals, to get initial values for *z_jl_* and *α*_*il*(*k*)_. To get the initial values of 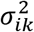, we apply DIMM-SC to each individual separately and get for each individual. Then we can match *α*_*i*(*k*)_ from the same cell type across different individuals based on calculating Pearson correlation and L1 norm of the difference between *α*_*i*(*k*)_ from different individuals. Specifically, each cell cluster was assigned to the cluster from another individual, which has the highest correlation with its gene expression profile and the lowest L1 norm of the difference between *α*_*i*(*k*)_ from different individuals.

### Section 2 Data generation in simulation studies

We simulated drop-seq data with different heterogeneity among multiple individuals. In our posited hierarchical model, the log normal prior distribution 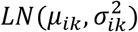 measures the heterogeneity of gene *i* in cell type *k* among multiple individuals and we also assume all 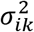’s follow a hyper-prior *Gamma*(*a_k_, b_k_*). For simplicity, we used the mean of 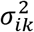 across all genes and all cell types to quantify the individual level heterogeneity.

In our simulation set-up, the UMI count matrix was sampled from a proposed Dirichlet mixture model. Specially, for a fixed total number of cell clusters *K,* we first pre-defined the values of *μ*_ik_, *a_k_* and *b_k_* for each cell cluster, and then sampled the 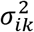 from the Gamma distribution *Gamma*(*a_k_, b_k_*). Next, we sampled *α*_*i.*(*k*)_ from a Log Normal distribution 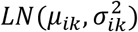 for each gene and each cell cluster. Proportion ***p_.jl_*** (*p*_1*jl*_, *p*_2*jl*_, …, *p*_*Gjl*_) was sampled from a Dirichlet distribution with parameter vector ***α*_*.l*(*k*)_** · (*α*_1*l*(*k*)_, *α*_2*l*(*k*)_, …, *α*_*Gl*(*k*)_, Last, we sampled the UMI count vector ***x*_*.jl*_** · (*x*_1*jl*_, *x*_2*jl*_, …, *x*_*Gjl*_ for the cell *j* and individual *l* from the multinomial distribution *Multi*(*T*_jl_, ***p*_*.jl*_**). We fixed 7}¡ as a constant across all cells and individuals.

### Section 3 Normalized mutual information

The mutual information (MI) of two random variables is a measure of the mutual dependence between the two variables. It measures the information these two variables share and how much knowing one of these variables reduces uncertainty about the other. Mutual information can be expressed as

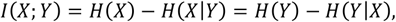

where *H*(*X*) and *H*(*Y*) are the marginal entropies, *H*(*X*|*Y*) and *H*(*Y*|*X*) are the conditional entropies. The normalized mutual information (NMI) can be expressed as

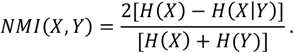

It represents the harmonic mean of 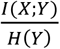 and 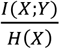 (Witten, et al., 2015). Since it’s normalized, we can measure and compare the NMI between different clustering results having different number of clusters.

---

**Supplementary Figure 1.**
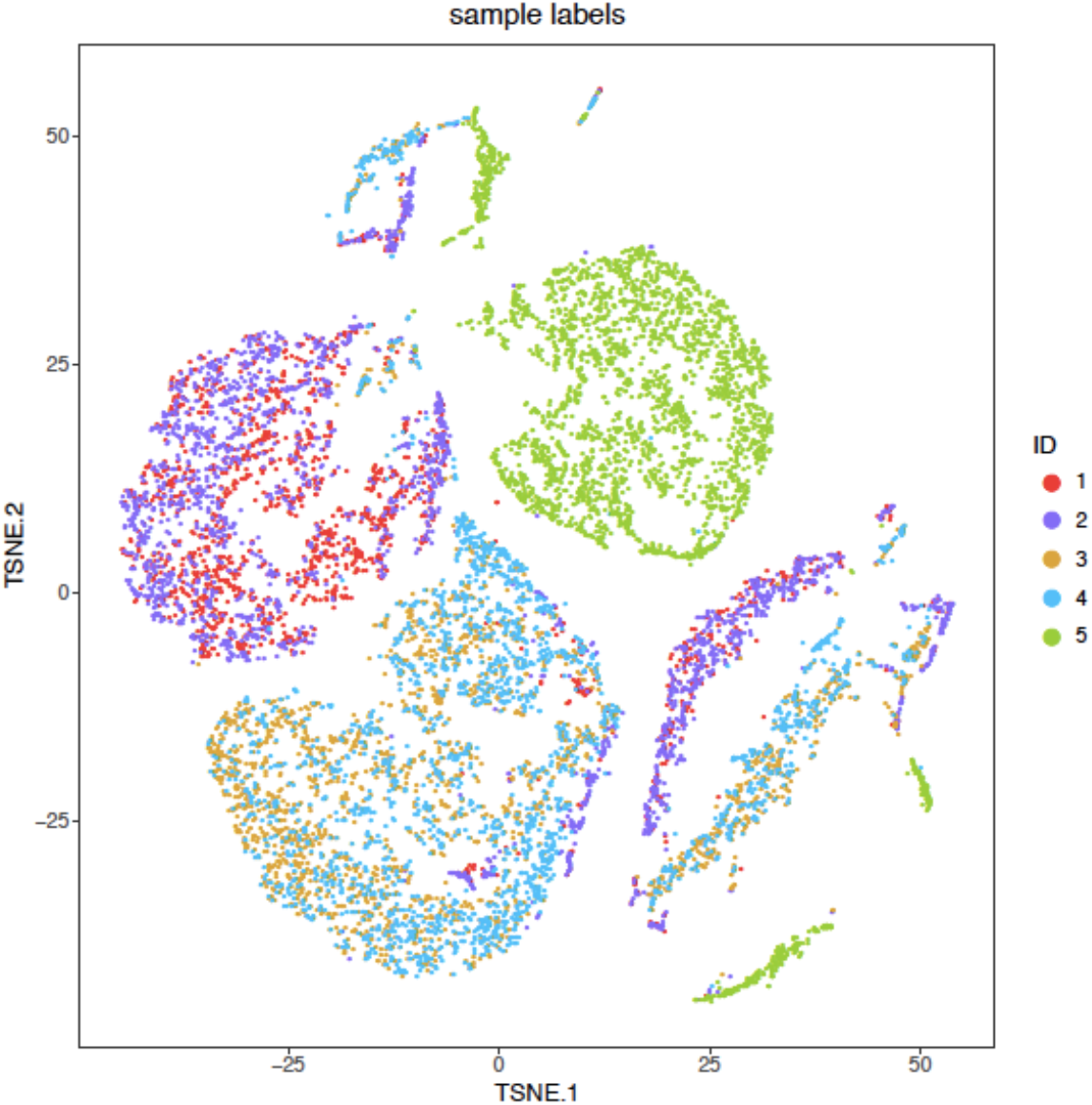
The t-SNE projection of human PBMC dataset, colored by different sample IDs.

**Supplementary Figure 2.**
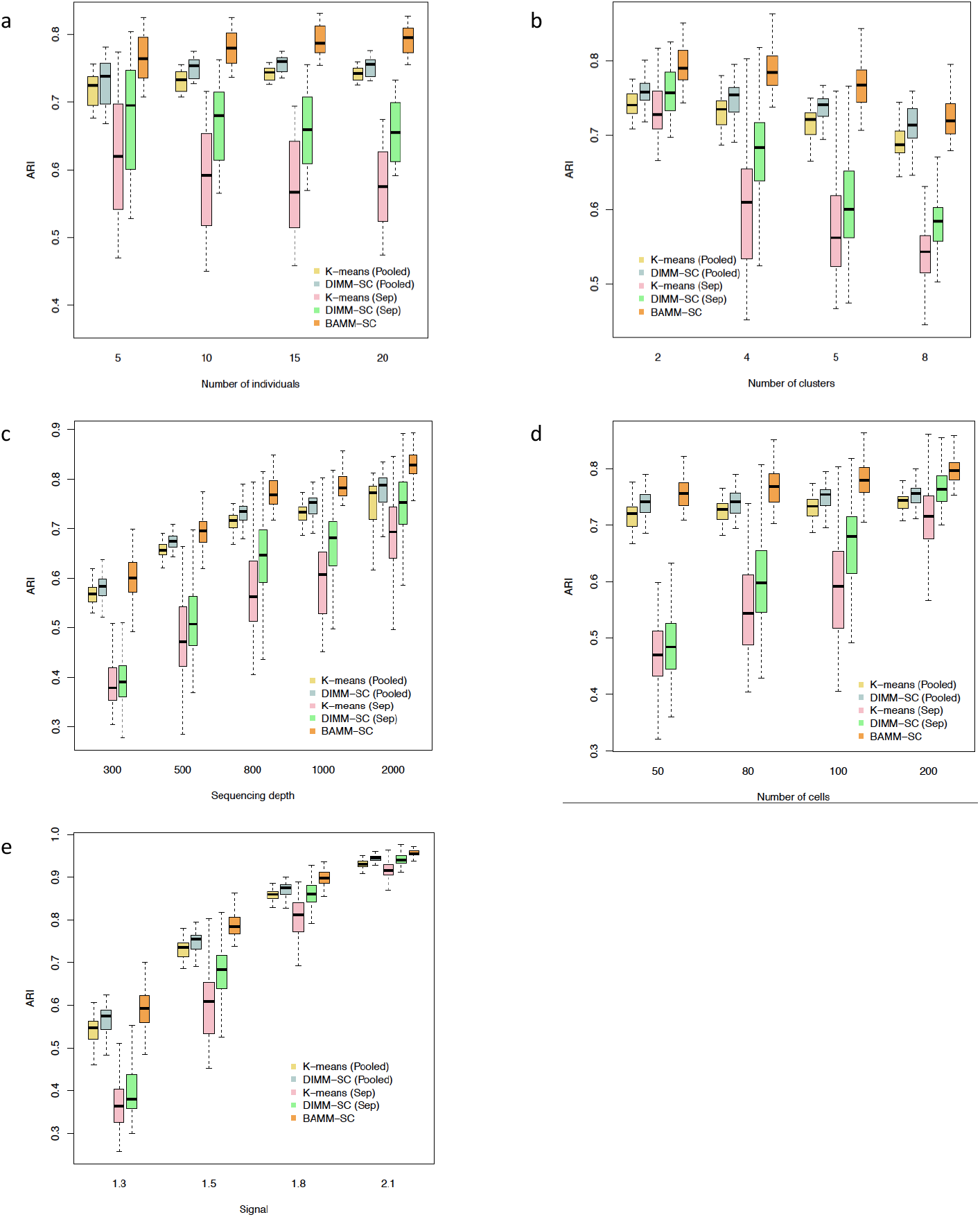
The Boxplots of ARI for five clustering methods across 100 simulations, investigating how different number of individuals (a), number of clusters (b), sequencing depth (c), number of cells in each cluster of each individual (d) and cell-type-specific heterogeneities (e) affect clustering results.

**Supplementary Figure 3.**
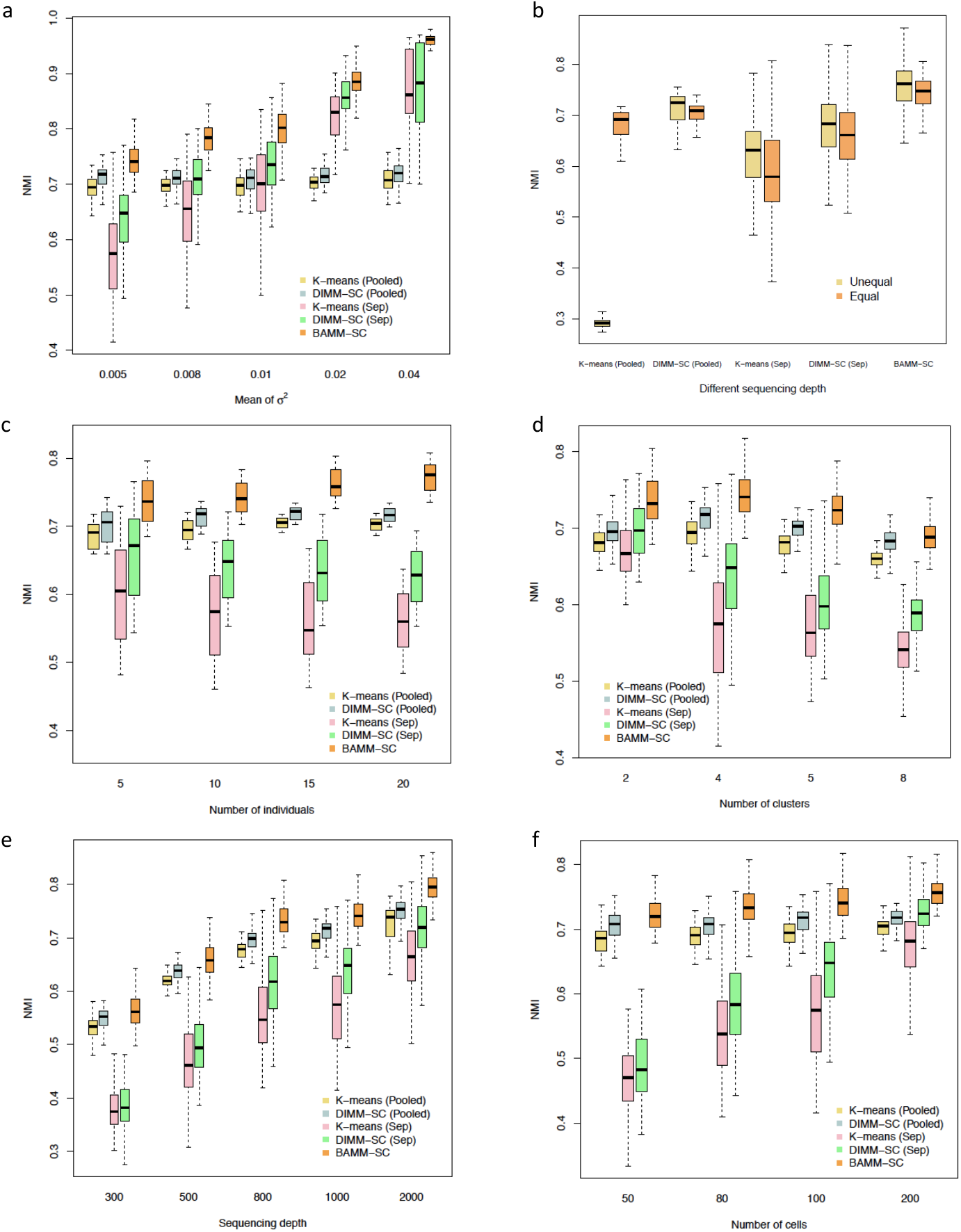
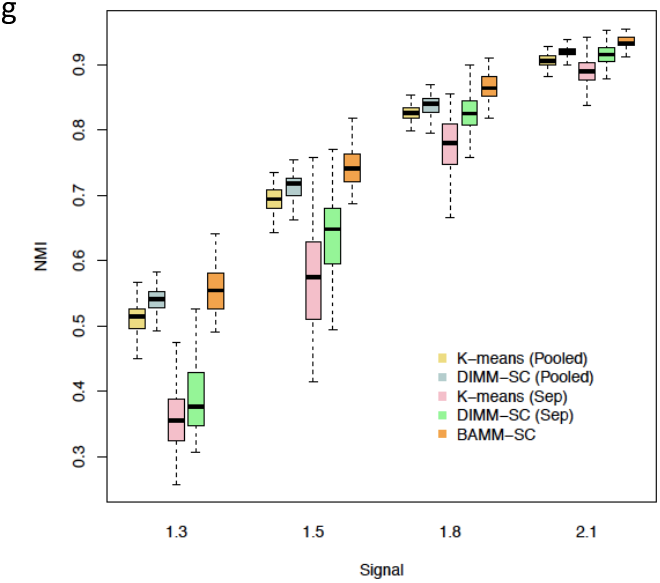
The Boxplots of NMI for five clustering methods across 100 simulations, investigating how different heterogeneity among multiple individuals (a), individuals with different sequencing depths (b), number of individuals (c), number of clusters (d), sequencing depth (e), number of cells in each cluster of each individual (f) and cell-type-specific heterogeneities (g) affect clustering results.

**Supplementary Figure 4.**
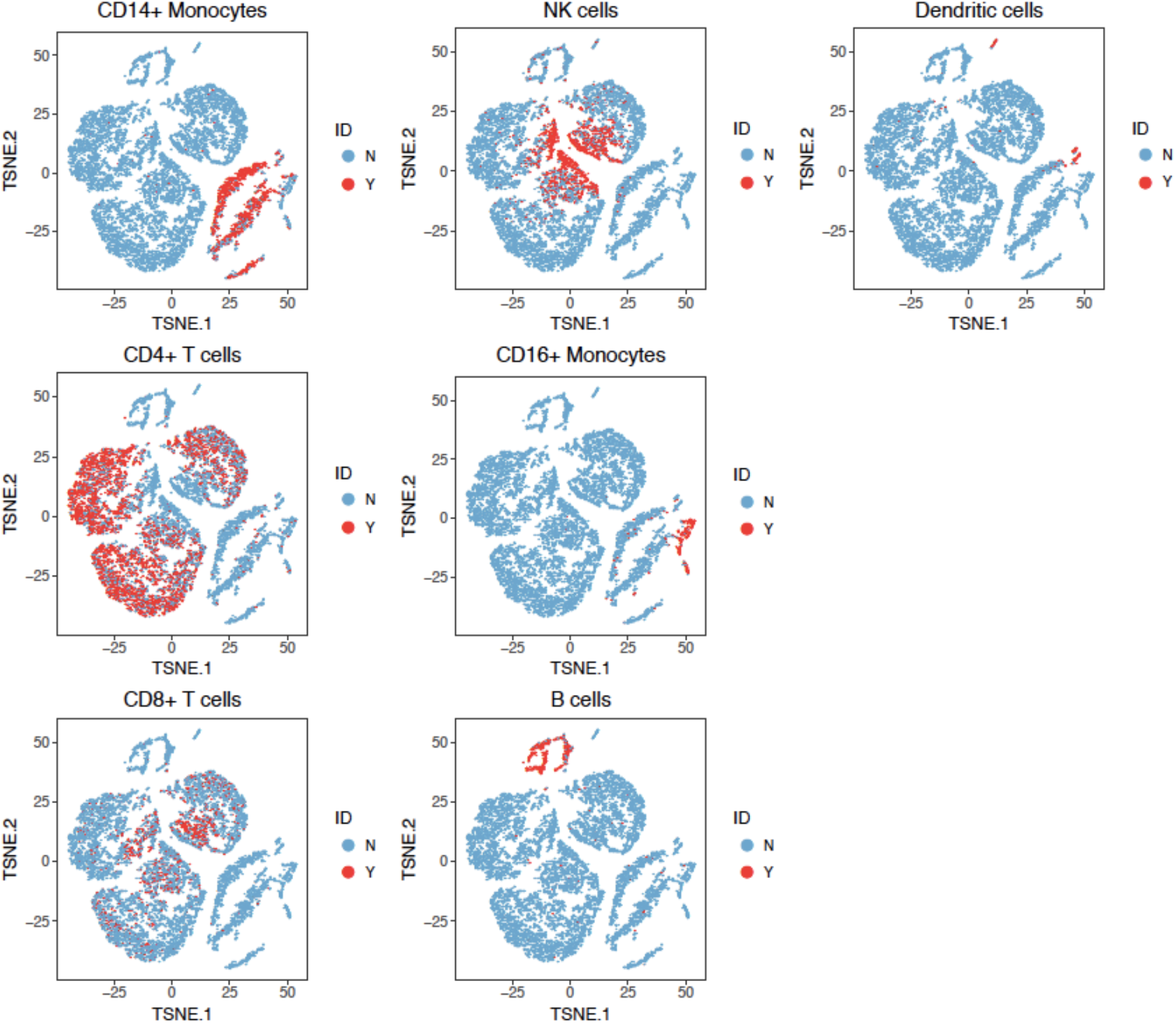
The t-SNE projection of cells from human PBMC, colored by different types of PBMCs based on the biological knowledge of cell-type-specific gene markers.

**Supplementary Figure 5.**
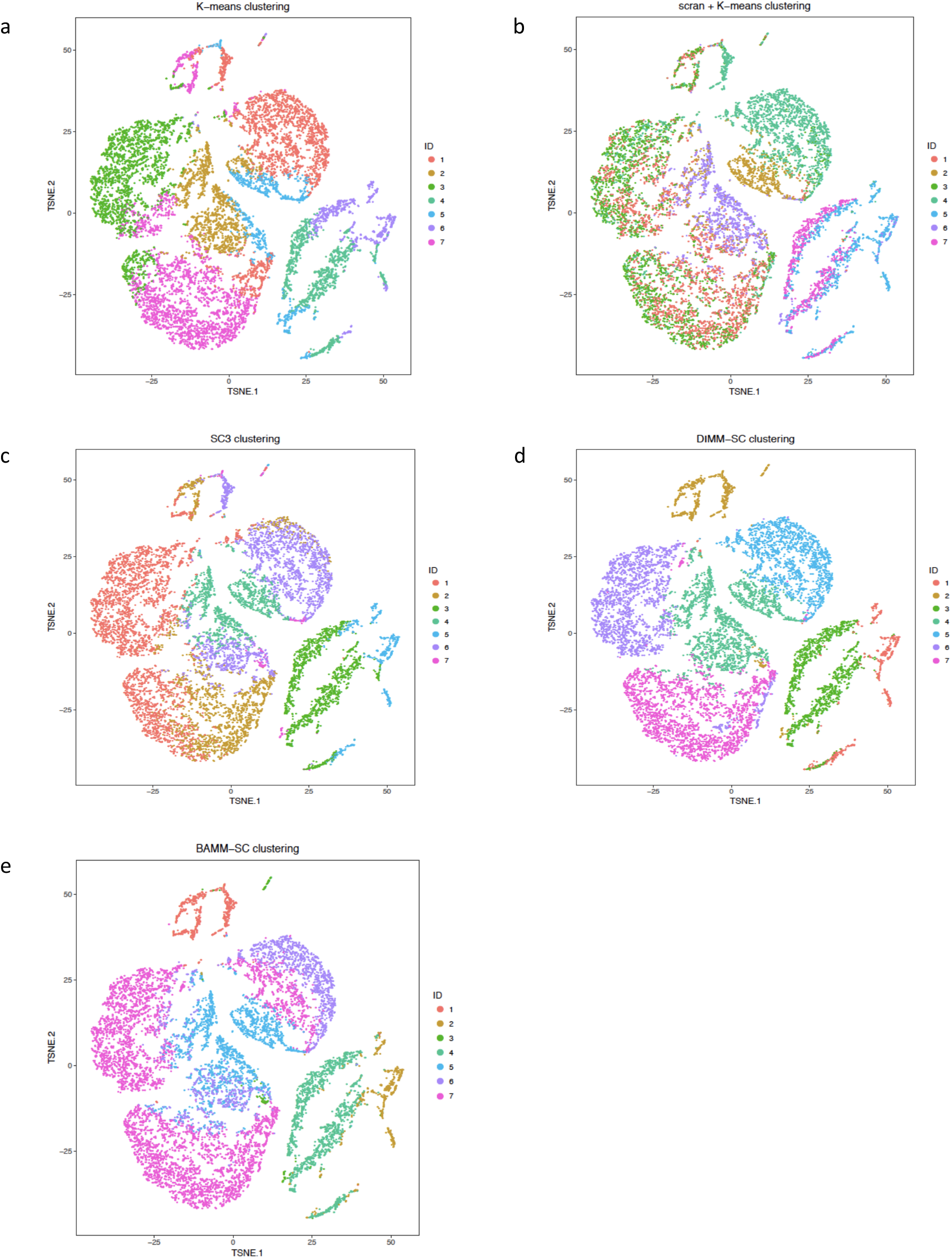
The t-SNE projection of cells from human PBMC, colored by the K-means clustering (a), scran + K-means clustering (b), SC3 clustering (c), DIMM-SC clustering (d) and BAMM-SC clustering assignment.

**Supplementary Figure 6.**
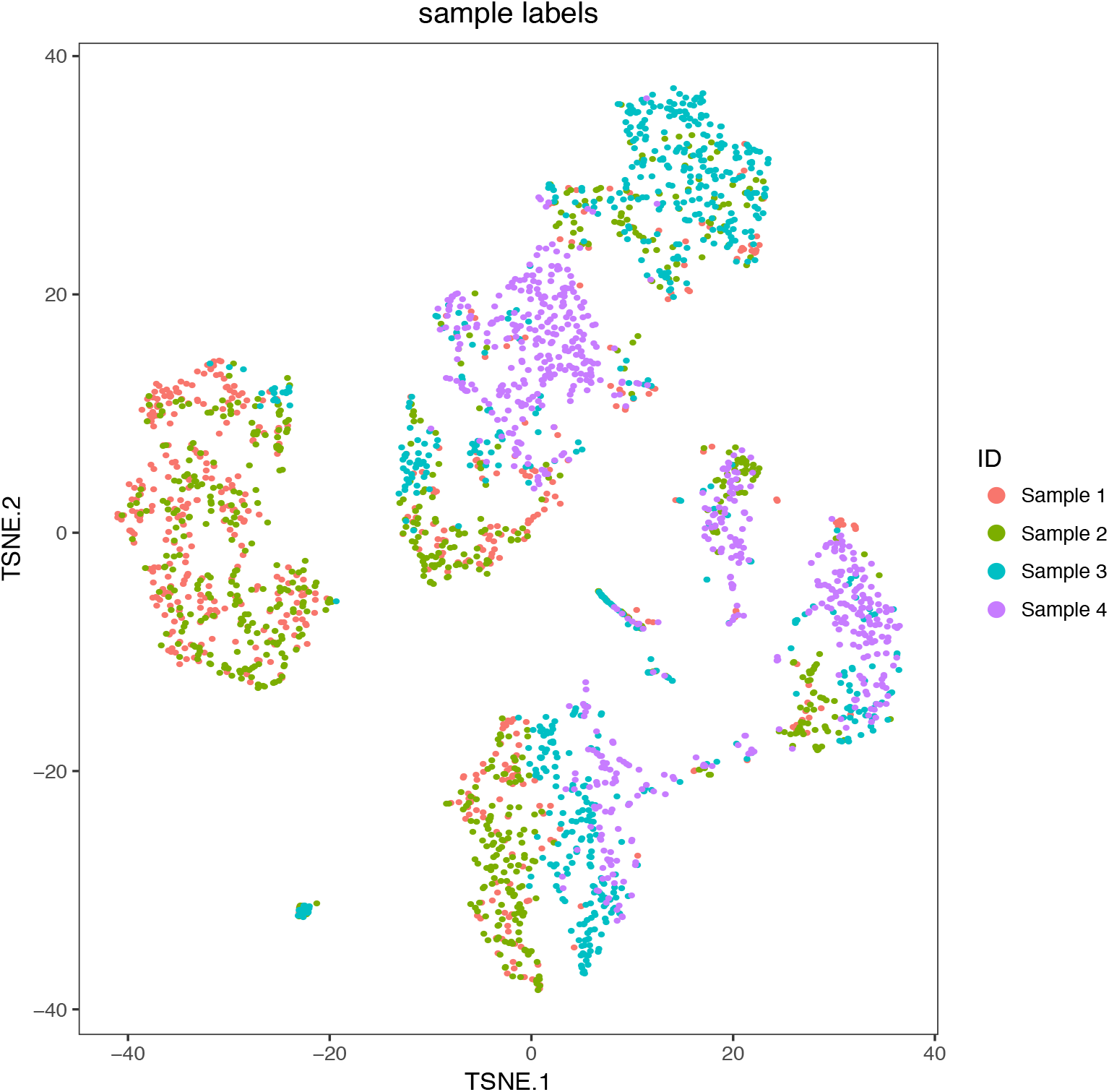
The t-SNE projection of mouse lung dataset, colored by different sample IDs.

**Supplementary Figure 7.**
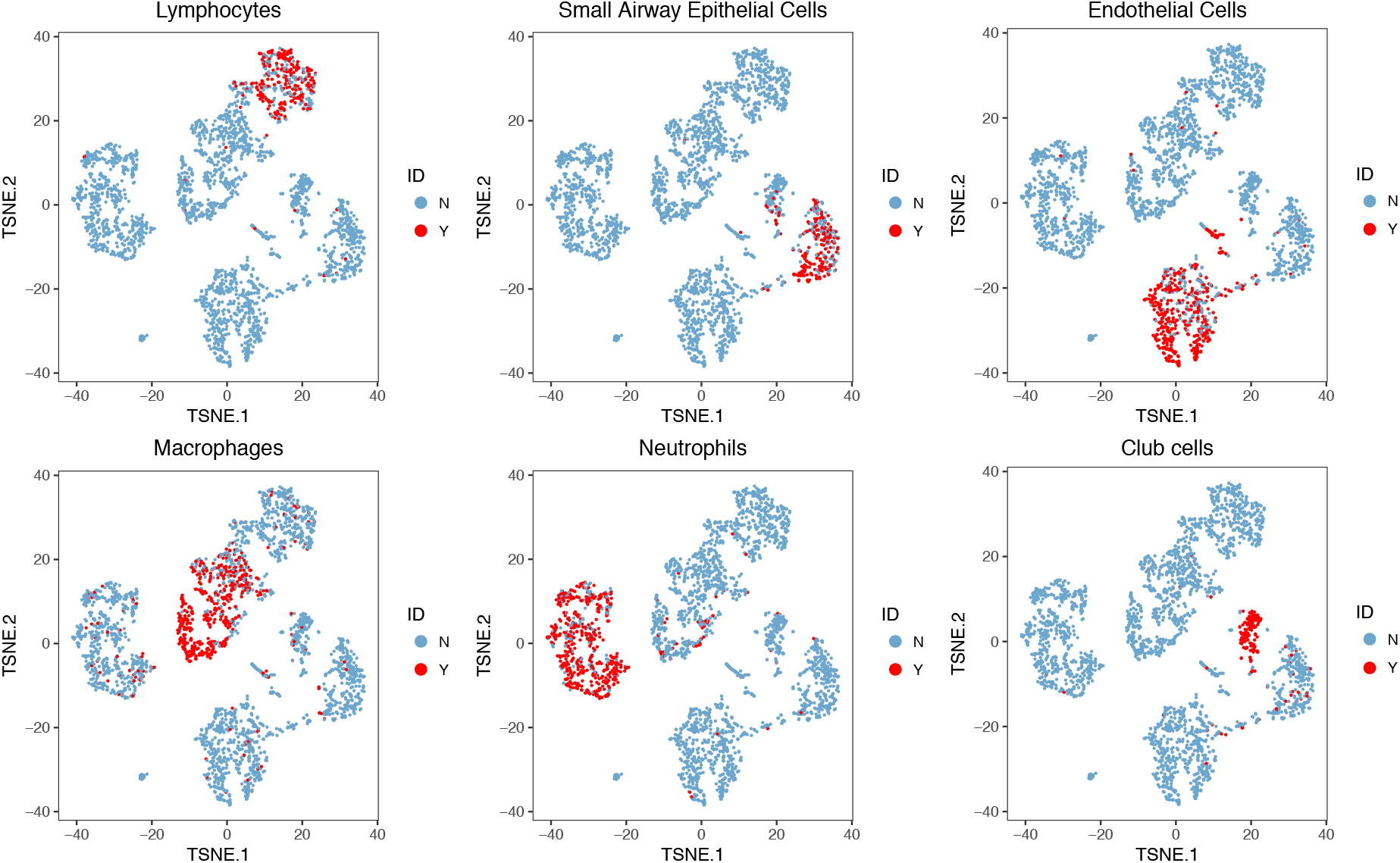
The t-SNE projection of cells from mouse lung dataset, colored by different types of PBMCs based on the biological knowledge of cell-type-specific gene markers.

**Supplementary Figure 8.**
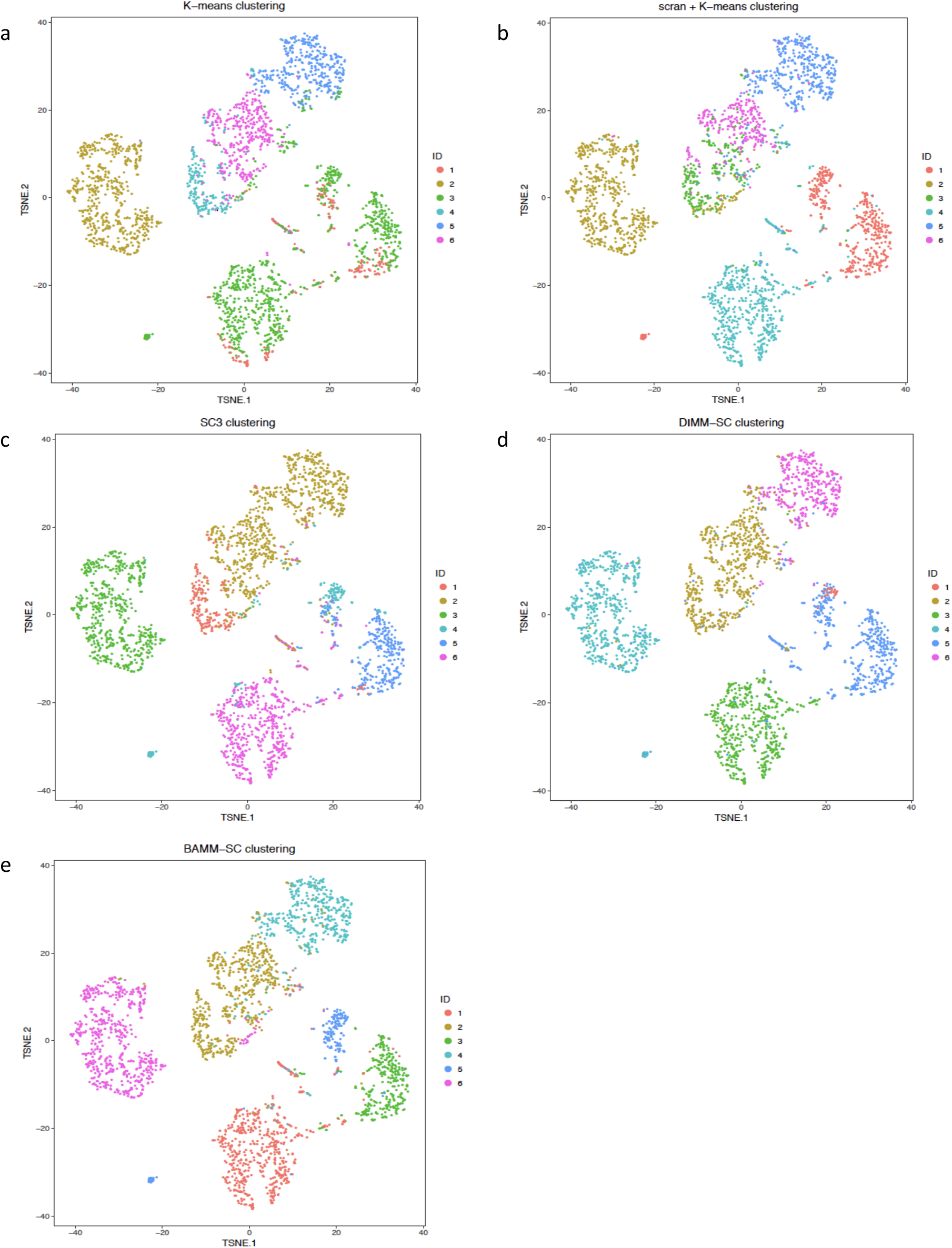
The t-SNE projection of cells from mouse lung dataset, colored by the K-means clustering (a), scran + K-means clustering (b), SC3 clustering (c), DIMM-SC clustering (d) and BAMM-SC clustering assignment.

**Supplementary Figure 9.**
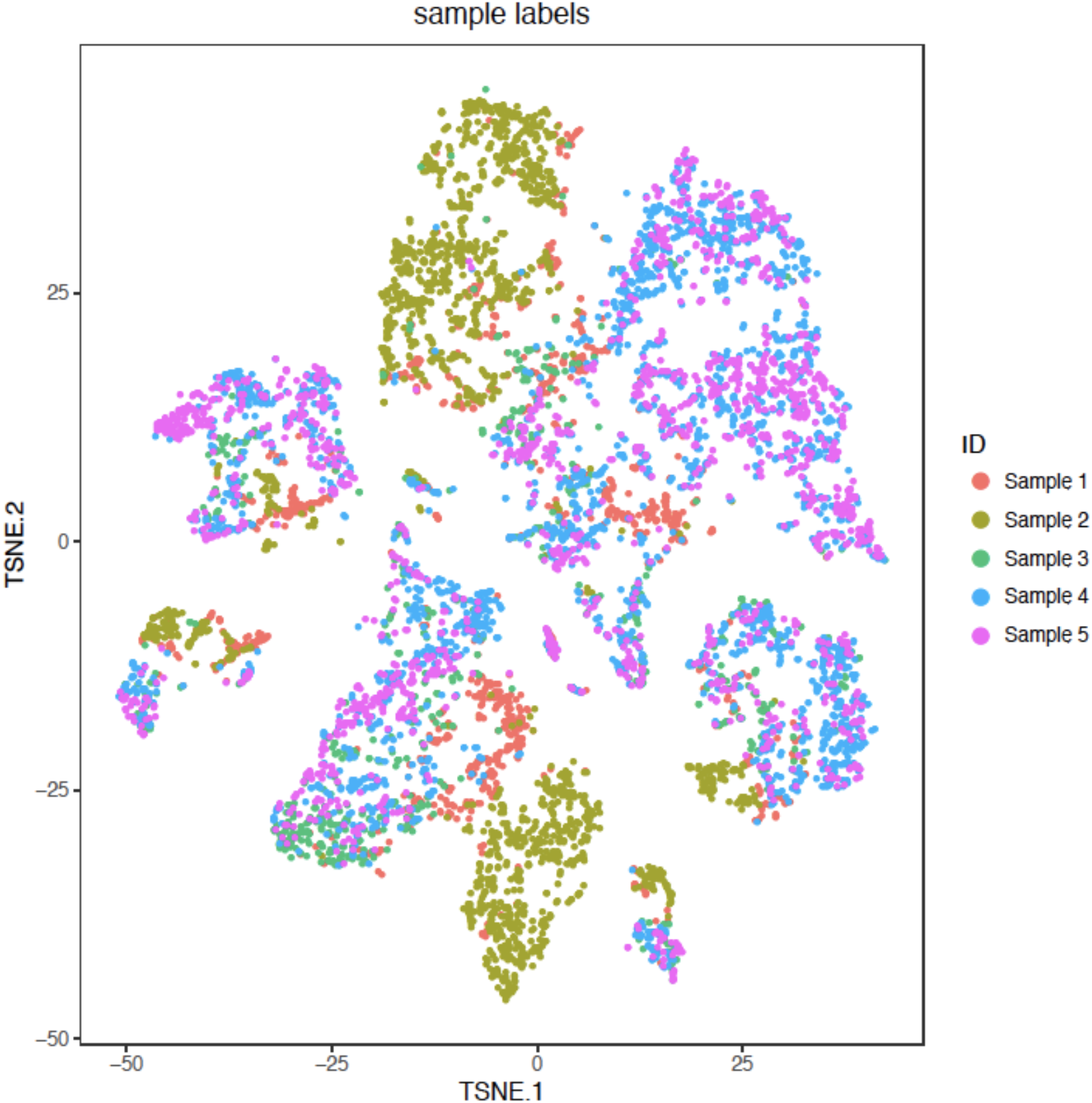
The t-SNE projection of cells from human skin dataset, colored by different sample IDs.

**Supplementary Figure 10.**
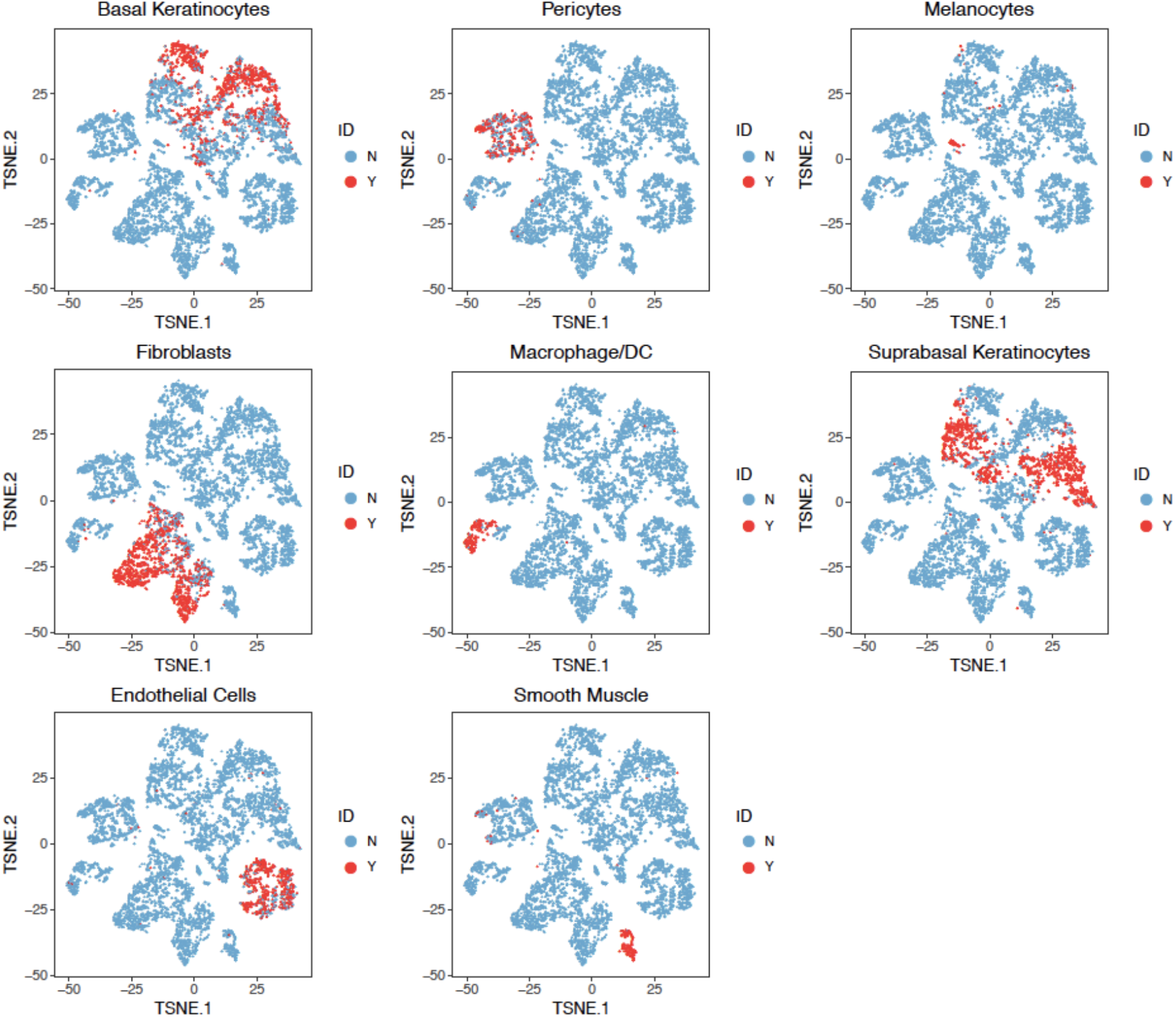
The t-SNE projection of cells from human skin dataset, colored by different types of PBMCs based on the biological knowledge of cell-type-specific gene markers.

**Supplementary Figure 11.**
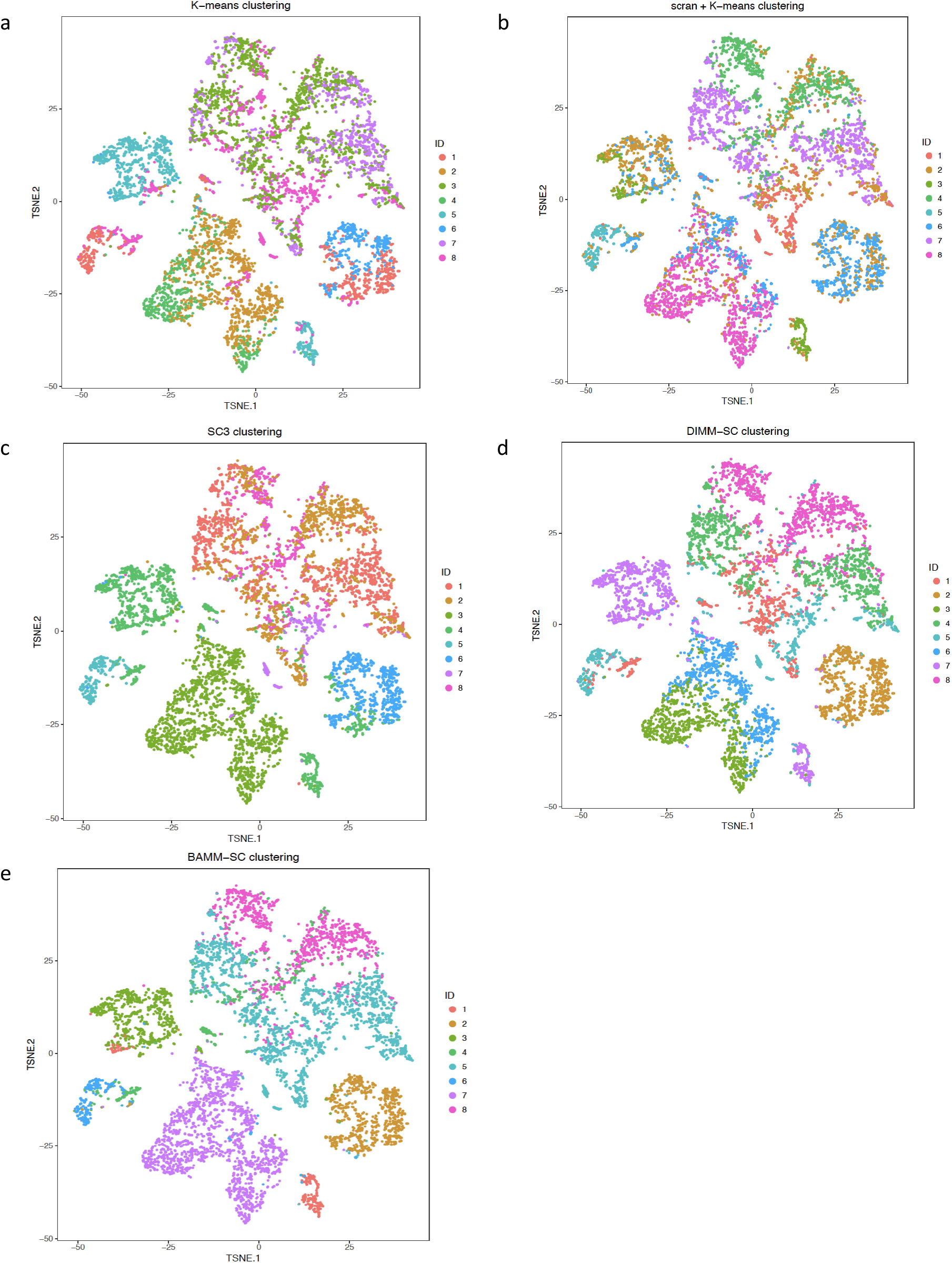
The t-SNE projection of cells from human skin dataset, colored by the K-means clustering (a), scran + K-means clustering (b), SC3 clustering (c), DIMM-SC clustering (d) and BAMM-SC clustering assignment.

**Supplementary Figure 12.**
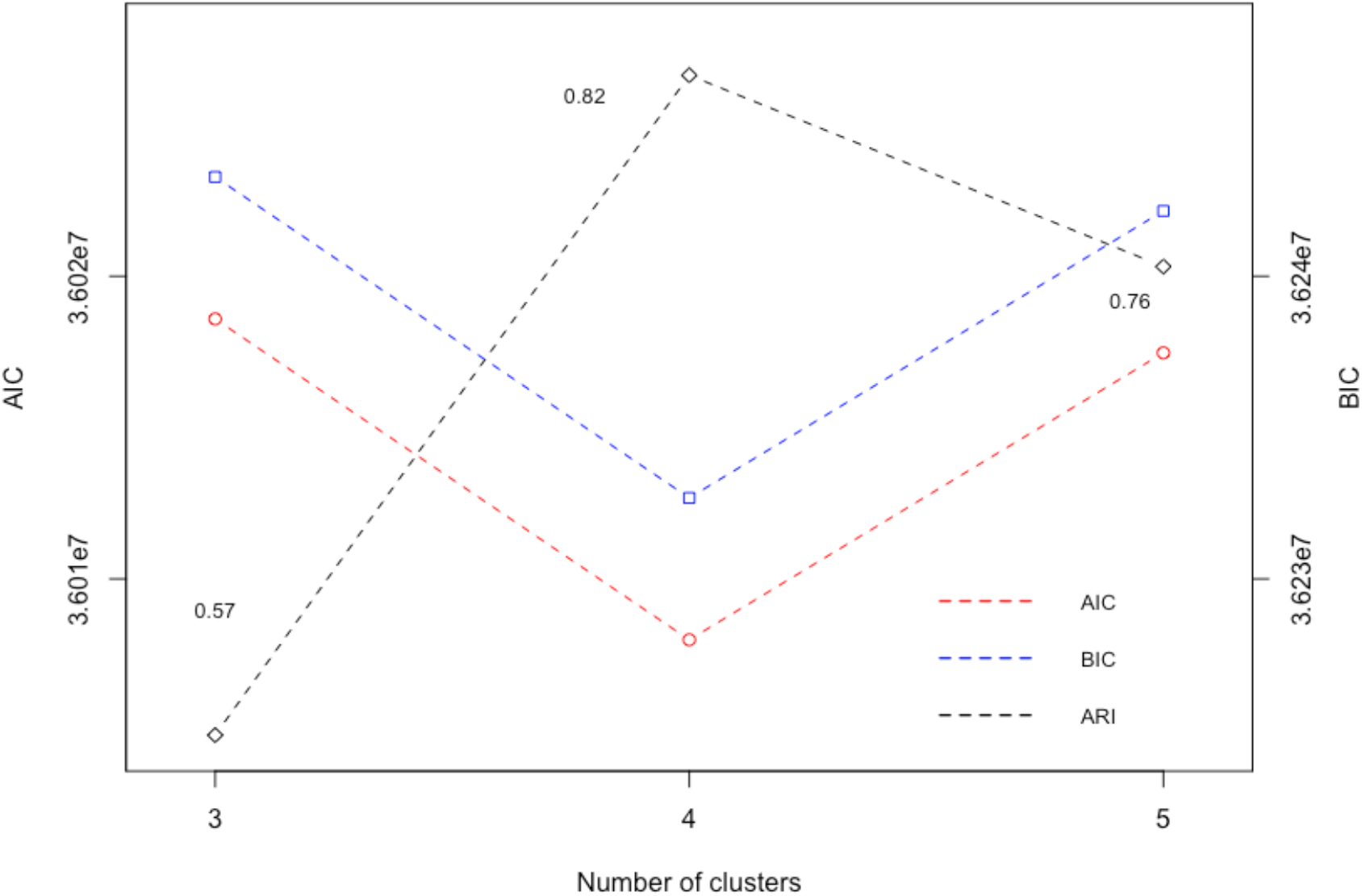
The dot plots of AIC and BIC for the final clustering results in the simulated dataset, where the true number of clusters is 4. Blue dots and red dots denote values of BIC and AIC, respectively. Black dots denote ARIs.

**Supplementary Figure 13.**
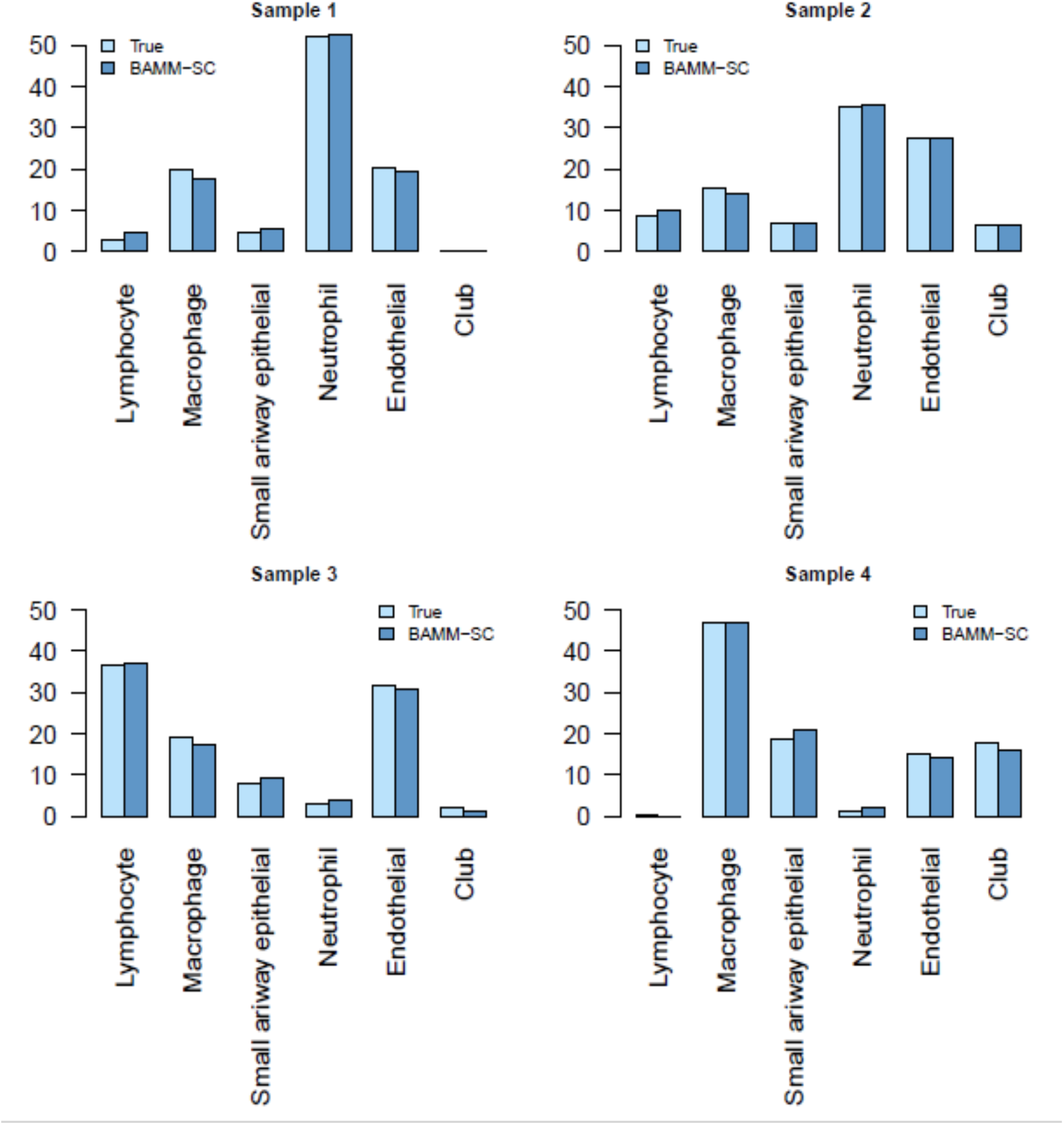
Bar plots of proportions of cell types for each individual in mouse lung dataset.

**Supplementary Table 1.**
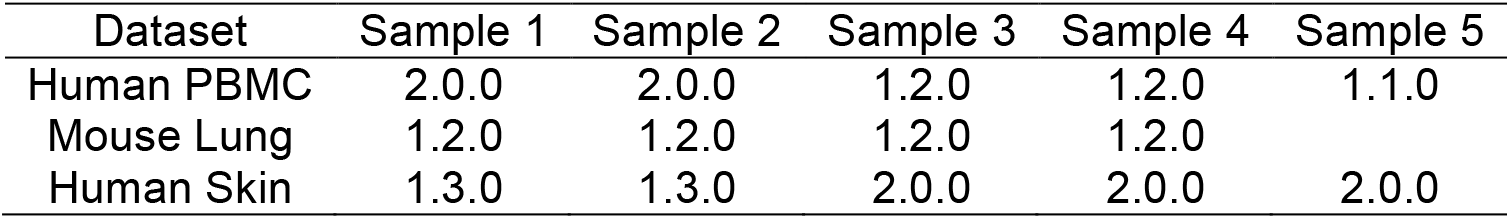
The version of Cell Ranger used when analyzing samples from three droplet-based scRNA-seq datasets.

**Supplementary Table 2.** Gene markers used to specify cell types in human PBMC samples.

| Cell Types | Genes |
| --- | --- |
| CD8+ T cells | CD3+CD8A+CD4- |
| CD4+ T cells | CD3+CD8-CD4+IL2RA-IL7R+ |
| B cells | CD3-CD19+MS4A1+ |
| NK cells | NCAM1+NKG7+CD3- |
| CD14+ monocytes | CD3-CD19-CD14+HLA- |
| CD16+ monocytes | CD3-CD19-FCGR3A+ |
| Dendritic cells | CD1C+CD14-HLA-FCER1A+ |

**Supplementary Table 6.**
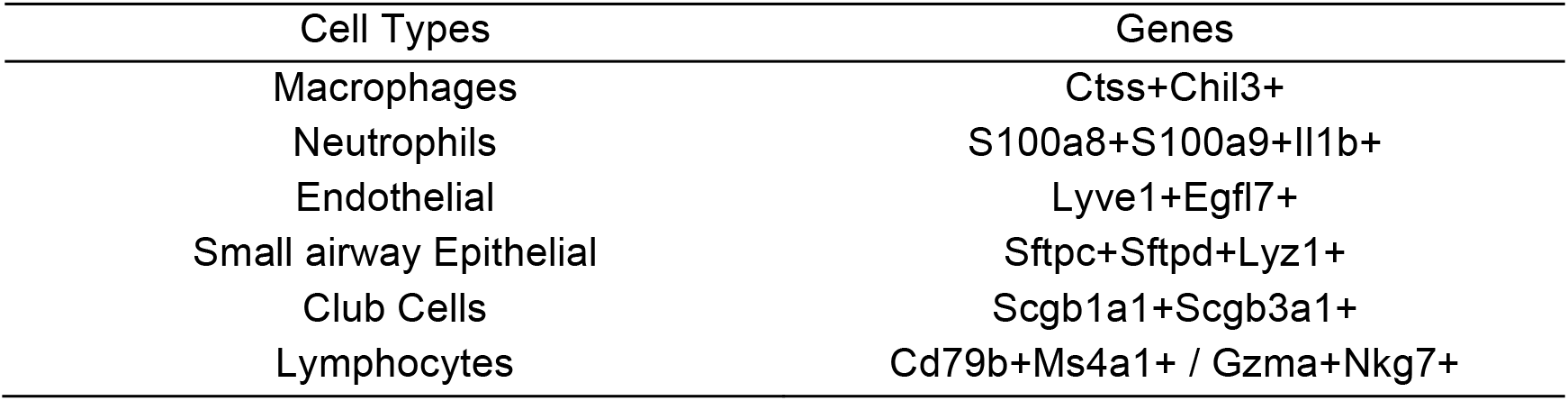
Gene markers used to specify cell types in mouse lung cell samples.

**Supplementary Table 4.** Gene markers used to specify cell types in human skin samples.

| Cell Types | Genes |
| --- | --- |
| Smooth muscle cells | DES+ |
| Suprabasal keratinocytes | KRT1+KRT10+ |
| Basal keratinocytes | KRT14+KRT5+ |
| Endothelial cells | VWF+ |
| Fibroblasts | COL1A1+ |
| Pericytes | RGS5+VWF- |
| Melanocytes | PMEL+ |
| Mecrophages/Dendritic cells | AIF1+ |

## References

1. Gawad, C., Koh, W. & Quake, S. R. Single-cell genome sequencing: current state of the science. Nat. Rev. Genet. 17, 175–188 (2016).

2. Tang, F. et al. mrna-seq whole-transcriptome analysis of a single cell. Nat. Methods 6, 377–382 (2009).

3. Macosko, E. Z. et al. Highly Parallel Genome-wide Expression Profiling of Individual Cells Using Nanoliter Droplets. Cell 161, 1202–1214 (2015).

4. Zheng, G. X. et al. Massively parallel digital transcriptional profiling of single cells. Nat. Commun. 8, 14049 (2017).

5. Jaitin, D. A. et al. Massively parallel single-cell RNA-seq for marker-free decomposition of tissues into cell types. Science 343, 776–779 (2014).

6. Pollen, A. A. et al. Low-coverage single-cell mRNA sequencing reveals cellular heterogeneity and activated signaling pathways in developing cerebral cortex. Nature biotechnology 32, 1053 (2014).

7. van der Wijst, M. G. et al. Single-cell RNA sequencing identifies celltype-specific cis-eQTLs and co-expression QTLs. Nature genetics 50, 493 (2018).

8. Rodriguez, A. & Laio, A. Clustering by fast search and find of density peaks. Science 344, 1492–1496 (2014).

9. Wang, B. et al. SIMLR: a tool for large-scale single-cell analysis by multi-kernel learning. Proteomics 18, 1700232 (2018).

10. duVerle, D. A., Yotsukura, S., Nomura, S., Aburatani, H. & Tsuda, K. CellTree: an R/bioconductor package to infer the hierarchical structure of cell populations from singlecell RNA-seq data. BMC Bioinformatics 17, 363 (2016).

11. Kiselev, V. Y. et al. SC3: consensus clustering of single-cell RNA-seq data. Nature methods 14, 483 (2017).

12. Sun, Z. et al. DIMM-SC: a Dirichlet mixture model for clustering droplet-based single cell transcriptomic data. Bioinformatics 34, 139–146 (2017).

13. Satija, R., Farrell, J. A. & Gennert, D. Spatial reconstruction of single-cell gene expression data. Nature biotechnology 33, 495–502 (2015).

14. Lun, A. T., McCarthy, D. J. & Marioni, J. C. A step-by-step workflow for low-level analysis of single-cell RNA-seq data with Bioconductor. F1000Research 5 (2016).

15. Rand, W. M. Objective criteria for the evaluation of clustering methods. Journal of the American Statistical association 66, 846–850 (1971).

16. Chen, K. & Kolls, J. K. T Cell–mediated host immune defenses in the lung. Annual review of immunology 31, 605–633 (2013).

17. Weiser, J. N. The pneumococcus: why a commensal misbehaves. Journal of molecular medicine 88, 97–102 (2010).

18. Tabib, T., Morse, C., Wang, T., Chen, W. & Lafyatis, R. SFRP2/DPP4 and FMO1/LSP1 define major fibroblast populations in human skin. Journal of Investigative Dermatology 138, 802–810 (2018).

19. Trapnell, C. et al. The dynamics and regulators of cell fate decisions are revealed by pseudotemporal ordering of single cells. Nature biotechnology 32, 381 (2014).

20. Trapnell, C. Defining cell types and states with single-cell genomics. Genome research 25, 1491–1498 (2015).

## References

1. Witten, I.H., et al. (2005) Data Mining: Practical Machine Learning Tools and Techniques, Morgan Kaufmann, Amsterdam. ISBN 978-0-12-374856-0.

